# Unveiling the A-to-I mRNA editing machinery and its regulation and evolution in fungi

**DOI:** 10.1101/2023.10.18.562923

**Authors:** Chanjing Feng, Kaiyun Xin, Yanfei Du, Jingwen Zou, Xiaoxing Xing, Qi Xiu, Yijie Zhang, Rui Zhang, Weiwei Huang, Qinhu Wang, Cong Jiang, Xiaojie Wang, Zhensheng Kang, Jin-Rong Xu, Huiquan Liu

**Author notes:** These authors contributed equally to this work (Chanjing Feng, Kaiyun Xin, Yanfei Du, Jingwen Zou).

## Abstract

A-to-I mRNA editing occurs during fungal sexual reproduction with an unknown mechanism. Here, we demonstrated that the eukaryotic tRNA-specific heterodimeric deaminase FgTad2-FgTad3, not typically associated with mRNA editing, is responsible for A-to-I mRNA editing in *Fusarium graminearum*. This editing capacity relies on the interaction between FgTad3 and a sexual stage-specific protein called Ame1. The interaction emerged in Sordariomycetes. Key residues involved in the interaction have been identified. Expression and activity of FgTad2-FgTad3 are regulated through alternative promoters, alternative translation initiation, and post-translational modifications. FgTad2-FgTad3-Ame1 efficiently edits target mRNAs in yeasts, bacteria, and human cells, with significant implications for developing base editors in therapy and agriculture. This study reveals mechanisms, regulation, and evolution of RNA editing in fungi, emphasizing protein-protein interactions in controlling enzyme function.

## Introduction

Adenosine-to-inosine (A-to-I) editing via deamination is a remarkable type of RNA editing as it modifies a multitude of nuclear-encoded mRNAs across the animal kingdom (*1*). As inosine is recognized as guanosine (G) by the translation machinery, A-to-I editing of coding regions can result in protein recoding. A-to-I editing also occurs at positions 34 (the wobble position of the anticodon) and 37 of tRNAs. While A^37^ editing has been found only in eukaryotes, A^34^ editing is conserved in both eukaryotes and bacteria (*2*). A-to-I mRNA editing in animals is catalyzed by enzymes of the Adenosine Deaminase Acting on RNA (ADAR) family that target specific double-stranded RNA (dsRNA) structures (*1*). The ADAR family originated in the last common ancestor of extant metazoans and is a metazoan innovation (*3*). All ADARs share a conserved C-terminal deaminase domain and a variable number of N-terminal dsRNA binding domains (dsRBDs) that mediate substrate recognition. The enzyme responsible for A-to-I tRNA editing is known as tRNA-specific Adenosine Deaminase (Tad) or Adenosine Deaminase Acting on tRNA (ADAT). Tad1/ADAT1 mediates A^37^ editing and shares a similar adenosine deaminase domain with ADARs but lacks dsRBDs (*2, 4*). In bacteria, homodimeric TadA deaminizes A^34^, while in eukaryotes, a heterodimer composed of the catalytic subunit Tad2/ADAT2 and the structural subunit Tad3/ADAT3 deaminizes it (*2, 5*). TadA, Tad2, and Tad3 all carry the catalytic motifs of cytidine/dCMP deaminases (CDAs), except that the essential glutamate (E) residue for proton shuttling in deamination is replaced by a catalytically inactive residue, usually valine (V), in Tad3 (*4*). Both subunits are essential for cell viability. Structural studies have shown that the C-terminal CDA domain of Tad3 interacts stably with Tad2 to form the catalytic domain of the heterodimeric enzyme, and the N-terminal Tad3 forms a tRNA binding domain that is flexibly linked to the catalytic domain (*5–7*).

Recently, A-to-I mRNA editing has been discovered in filamentous ascomycetes, specifically during sexual reproduction (*8, 9*). It appears to be a common feature in Sordariomycetes (*10*), one of the largest classes of Ascomycota. More than 40,000 A-to-I mRNA editing sites have been identified in the perithecia (sexual fruiting bodies) of *Neurospora crassa* and *Fusarium graminearum*, respectively (*11, 12*). As fungi lack orthologs of animal ADARs and the *cis*-regulatory elements of editing are also distinct from animals (*12*), a different A-to-I editing mechanism must exist in fungi. Here, we demonstrated that FgTad2-FgTad3 is responsible for A-to-I mRNA editing in *F. graminearum,* with its capacity relying on the interaction between FgTad3 and a sexual stage-specific protein named Ame1 (<u>a</u>ctivator of <u>m</u>RNA <u>e</u>diting). This interaction emerged in the last common ancestor of Sordariomycetes, with key residues identified. Moreover, the FgTad2-FgTad3-Ame1 complex efficiently edits target mRNAs not only in yeasts and bacteria but also in human cell lines, advancing the development of new therapeutic and agricultural tools. Our study uncovers the mechanisms, regulation, and evolution of RNA editing in fungi, emphasizing the significance of protein-protein interactions in controlling deaminase function.

## Results

### FgTad2 and FgTad3 form a heterodimer in *F. graminearum*

Both FgTad2 and FgTad3 contain the typical CDA domain with conserved residues required for activities in their catalytic core, except that the conserved E residue essential for catalysis was replaced by a V residue in FgTad3 (Fig. 1A). In comparison with the orthologs in yeast, plants, and animals, the sequence upstream from the CDA domain of FgTad2 is unusually long. Besides the CDA domain (FgTad3^CDA^), FgTad3 also contains an N-terminal domain (FgTad3^N^) known for tRNA binding. During sexual development from 1- to 8-days post-fertilization (dpf), *FgTAD2* and *FgTAD3* showed similar expression levels and trends (Fig. 1B). Both yeast two-hybrid (Y2H) and co-immunoprecipitation (co-IP) assays confirmed their interaction (Fig. 1C-1D). It was found that FgTad2 interacts with the C-terminal (203-428 aa) but not the N-terminal (1-202 aa) part of FgTad3. Therefore, FgTad2 and FgTad3 function as a heterodimer in *F. graminearum* for adenosine deamination. Like their yeast orthologs, FgTad2 serves as the catalytic subunit, while FgTad3 plays a structural role.

**Fig. 1.**
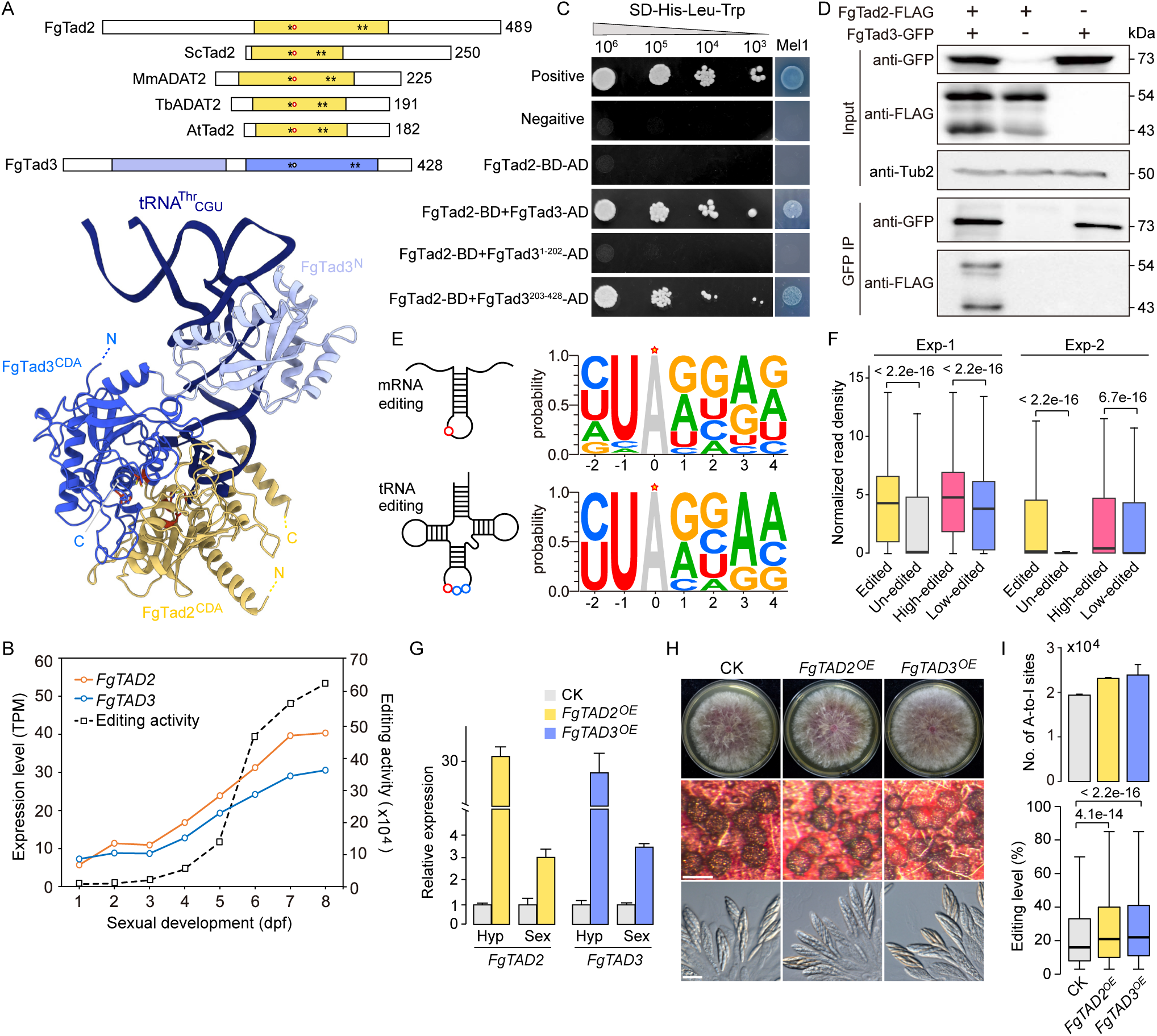
A-to-I editing of tRNA and mRNA by the FgTad2-FgTad3 complex. (A) Schematic representation of the domain structure of FgTad2 and FgTad3, along with Tad2/ADAT2 from *Saccharomyces cerevisiae* (Sc), *Mus musculus* (Mm), *Trypanosoma brucei* (Tb), and *Arabidopsis thaliana* (At). The AlphaFold model of the FgTad2-FgTad3 heterodimer bound to tRNA is also depicted. Tad2 CDA is represented in orange, FgTad3^CDA^ in blue, and FgTad3 N-terminal domain (FgTad3^N^) in sky blue. Asterisks (*) indicate Zn^2+^ coordinating residues. The active-site glutamate/pseudo-active-site valine are denoted by red and black circles, respectively. (B) Expression levels of *FgTAD2* and *FgTAD3*, as well as A-to-I mRNA editing activity (summing the editing levels of all editing sites) during sexual development from 1 to 8 dpf. (C) Yeast two-hybrid assays to detect the interaction between FgTad2 (BD-bait) and the full-length protein, N-terminal (1-202 aa), or C-terminal (203-428 aa) region of FgTad3 (AD-prey). Photos are presented showing growth on SD-His-Leu-Trp plates at indicated concentrations, along with the Mel1 α-galactosidase activity. (D) Co-immunoprecipitation (co-IP) assays for the interaction between FgTad2 and FgTad3. Western blots of total proteins isolated from transformants expressing the FgTad2-3×FLAG and/or FgTad3-GFP (input) and proteins eluted from anti-GFP affinity beads (GFP IP) were detected with anti-FLAG and anti-GFP antibodies. (E) Structures and sequence motifs of A-to-I mRNA editing sites and inosine-modified tRNA A^34^ sites in *F. graminearum*. Red circles and stars mark the editing site in the RNA structure diagram and sequence logo, respectively. (F) Boxplots of normalized read densities in the indicated gene regions from two independent experiments (Exp-1 and Exp-2) of FgTad2-FLAG RIP-Seq. *P* values are from the two-tailed Wilcoxon rank sum test. (G) qRT-PCR analysis of *FgTAD2* and *FgTAD3* expression levels in hyphae (Hyp) and perithecia (Sex) of the control check (CK) strain and overexpressing (OE) transformants. Mean and SD were calculated with data from three independent repeats (n = 3). (H) Colony morphology, perithecium formation, and asci/ascospores morphology of marked strains. Bar = 0.2 mm (Upper); bar = 20 μm (Bottom). (I) Number of A-to-I mRNA editing sites and editing levels in 7-dpf perithecia of marked strains. For the bar graph, Mean and SD were calculated with data from two independent repeats (n = 2). *P* values are from the two-tailed Wilcoxon rank sum test.

### FgTad2-FgTad3 binds to mRNA and prefers highly edited transcripts

The expression levels of *FgTAD2* and *FgTAD3* were found to be correlated with the activity of A-to-I mRNA editing during sexual development (Fig. 1B). In addition to being located within a similar stem-loop structure, A-to-I mRNA editing sites also share similar sequence preferences with the A^34^ sites of eight inosine-modified tRNAs in *F. graminearum* (Fig. 1E). To investigate the mRNA-binding capability of the FgTad2-FgTad3 complex, we generated transformants *in situ* expressing a *FgTAD2*-FLAG construct and performed RNA immunoprecipitation sequencing (RIP-seq) with 7-dpf perithecia using an anti-FLAG antibody. The RIP-seq experiments were repeated twice with independent biological replicates. In both experiments, the density of footprint reads was high in edited transcripts but low in unedited transcripts expressed in perithecia (Fig. 1F). Moreover, highly-edited transcripts had significantly higher footprint read densities relative to low-edited transcripts. These data indicate that FgTad2-FgTad3 has an mRNA-binding capacity and that the mRNA-binding capacity is correlated with the editing intensity of transcripts. Therefore, FgTad2-FgTad3 is also responsible for mRNA editing in *F. graminearum*.

### Overexpression of *FgTAD2* or *FgTAD3* enhances mRNA editing activities in sexual reproduction

To confirm the function of FgTad2-FgTad3 in mRNA editing, overexpression analysis was performed since deletion of these genes is lethal. An extra copy of *FgTAD2* or *FgTAD3* was introduced in PH-1 at a targeted locus under the control of the constitutive RP27 promoter. Real-time quantitative Reverse Transcription PCR (qRT-PCR) analysis showed that the expression levels of *FgTAD2* and *FgTAD3* in the resulting transformants were increased over 20-fold in 24-h vegetative hyphae and 3-fold in 7-dpf perithecia (Fig. 1G). Transformants overexpressing *FgTAD2* or *FgTAD3* showed normal vegetative growth and only a minor defect in sexual development. In both transformants, perithecia were slightly transparent in color (Fig. 1H). Strand-specific RNA-seq analysis revealed that the number of A-to-I mRNA editing sites and their median editing levels were increased in 7-dpf perithecia of both transformants (Fig. 1I). These results indicate that increasing the expression of either FgTad2 or FgTad3 can enhance mRNA editing activities in sexual reproduction.

### Alternative promoter usage results in the expression of a short transcript isoform of both *FgTAD2* and *FgTAD3* during sexual reproduction

Interestingly, RNA-seq data showed that both *FgTAD2* and *FgTAD3* expressed two major transcript isoforms via alternative transcriptional initiation (Fig. 2A). The long (L) transcript was constitutively expressed, while the short (S) transcript was expressed specifically during the sexual stage. For both genes, the abundance of the L-transcript remained relatively constant during sexual development, while the S-transcript was continuously upregulated after 4-dpf and became the dominant transcript in the late stage of sexual development (Fig. 2B). These data suggest that the S-transcript is the main mediator for mRNA editing.

**Fig. 2.**
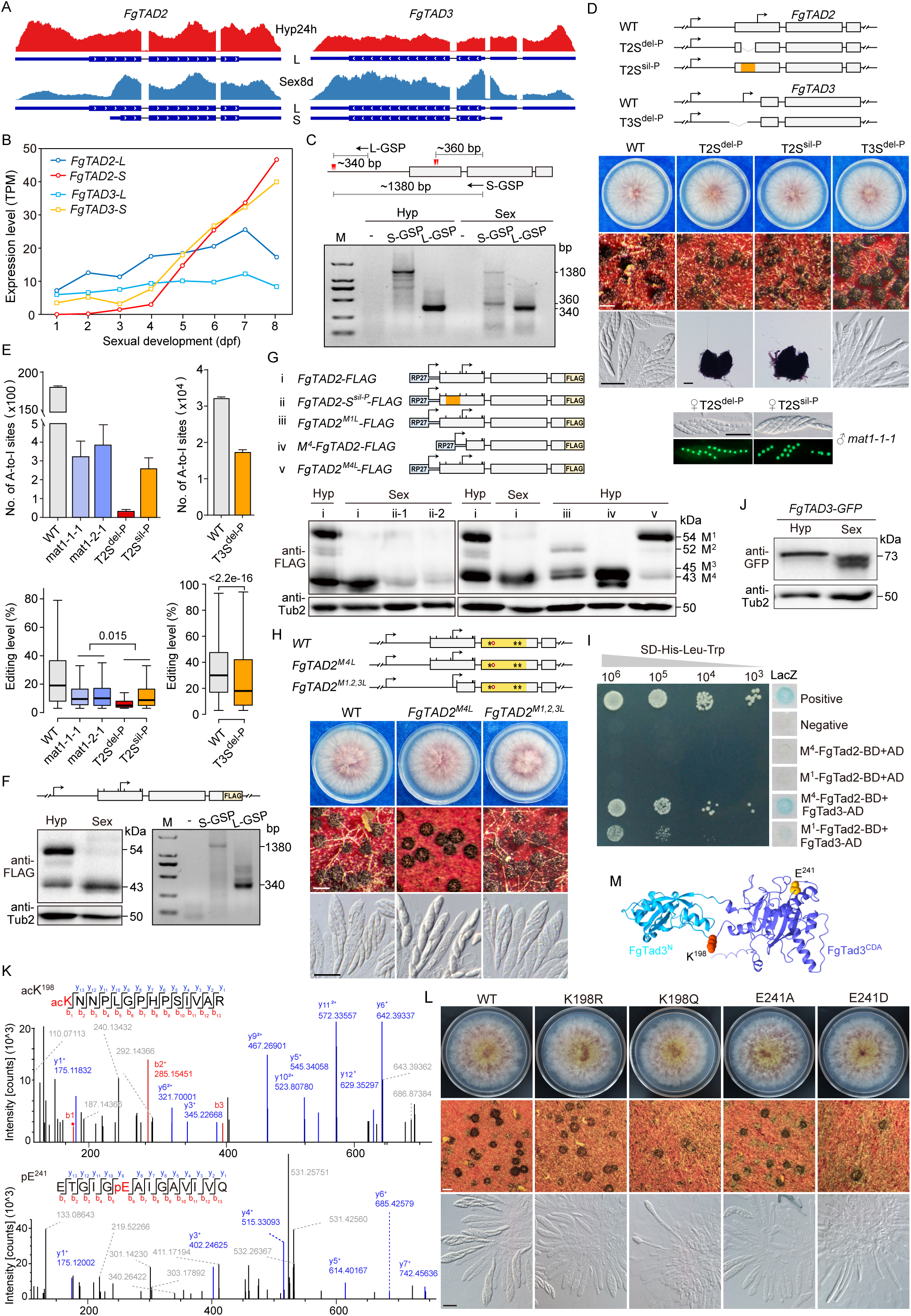
Transcriptional, translational, and post-translational regulation of *FgTAD2* and *FgTAD3*. (A) RNA-Seq read coverage showing the expression of long (L) and short (S) transcript isoforms of *FgTAD2* and *FgTAD3* during vegetative growth (Hyp24h) and sexual reproduction (Sex8d). (B) Expression levels of the L and S transcripts of *FgTAD2* and *FgTAD3* during sexual development from 1 to 8 dpf. (C) 5’RACE determining the transcriptional initiation sites (marked by red triangles) of the two *FgTAD2* transcripts. The expected size for each band is indicated. M, DNA markers; GSP, gene-specific primers. (D) Colony morphology, perithecium formation, and asci/ascospores morphology of the mutants with deletion of the promoter region of the *FgTAD2* S-transcript (T2S^del-P^) or *FgTAD3* S-transcript (T3S^del-P^) and the mutant with multiple synonymous mutations introduced in the promoter region of the *FgTAD2* S-transcript (T2S^sil-P^). Morphology of ascospores from crosses of the marked mutants (♀) with the *mat1-1-1* mutant labeled with an H1-GFP (♂) are also shown. White bar = 0.2 mm; black bar = 20 μm. In the schematic diagram, arrows indicate transcriptional initiation sites, and boxes indicate CDS regions. Broken lines represent deletion regions, while the region that introduced multiple synonymous mutations is highlighted in orange. (E) Number of A-to-I mRNA editing sites detected in 7-dpf perithecia of marked strains, as well as the editing levels of common editing sites. For the bar graph, Mean and SD were calculated with data from two independent repeats (n = 2). *P* values are from the two-tailed Wilcoxon rank sum test. (F) Western blots of total proteins isolated from 24-h hyphae (Hyp) and 7-dpf perithecia (Sex) of the transformant *in situ* expressing the FgTad2-3×FLAG, probed with anti-FLAG antibodies. The 5’RACE analysis confirmed that only the *FgTAD2* L-transcript was expressed in the hyphae of this transformant. (G) Western blots of total proteins isolated from 24-h hyphae (Hyp) and 7-dpf perithecia (Sex) of transformants ectopically expressing the indicated constructs. The location of first seven in-frame AUG (M) codons are indicated with a bar on the CDS box. (H) Colony morphology, perithecium formation, and asci/ascospores morphology of the marked *FgTAD2* mutants. White bar = 0.2 mm; black bar = 20 μm. The CDA domain region is highlighted in orange in the schematic diagram. (I) Yeast two-hybrid assays for the interaction of the FgTad2 M^1^- or M^4^-protein (BD-bait) with FgTad3 (AD-prey). Photos are presented showing growth on SD-His-Leu-Trp plates at indicated concentrations, along with the LacZ β-galactosidase activity. (J) Western blots of total proteins isolated from 24-h hyphae (Hyp) and 7-dpf perithecia (Sex) of the transformant ectopically expressing the FgTad3-GFP, probed with anti-GFP antibodies. (K) MS/MS spectrum of acetylated and phosphorylated peptides. acK^198^ and pE^241^ indicate the location of acetylation and phosphorylation, respectively. (L) Colony morphology, perithecium formation, and asci/ascospores morphology of the modification-mimetic and -deficient mutants for the acK^198^ and pE^241^ of *FgTAD3*. White bar = 0.2 mm; black bar = 20 μm. (M) Cartoon model depicting the location of K^198^ and E^241^ sites in FgTad3.

For *FgTAD2*, the transcriptional initiation site of the S-transcript was located within the open reading frame (ORF) region of the L-transcript according to the RNA-seq data (Fig. 2A). We used the 5’RACE (Rapid Amplification of 5’-Ends cDNA) method to determine the precise transcriptional initiation sites of the two *FgTAD2* transcript isoforms. A PCR band with a size of ∼340 bp associated with the L-transcript initiation was detected in both hyphae and perithecia, while a band with a size of ∼360 bp associated with the S-transcript initiation was detected only in perithecia (Fig. 2C), confirming the unique transcription of the S-transcript during sexual reproduction. Sequencing of these PCR products revealed that the transcriptional initiation sites of the L-transcript were located between −743 and −713 bp upstream of the start codon, and those of the S-transcript were located between 280-310 bp downstream of the start codon of the L-transcript.

### The *FgTAD2* S-transcript plays a critical role in mRNA editing while the *FgTAD3* S-transcript is important for mRNA editing

To verify the role of the *FgTAD2* S-transcript in sexual reproduction and mRNA editing, we removed its putative promoter region (85-264 bp relative to the start codon of the L-transcript) at the native locus. The resulting T2S^del-P^ mutant was normal in vegetative growth but produced smaller perithecia without asci and ascospores (Fig. 2D). We also generated mutants with multiple synonymous mutations introduced in the promoter region of the S-transcript, which aimed to suppress the transcription of the S-transcript without affecting the protein encoded by the L-transcript. The resulting T2S^sil-P^ mutant had similar defects as the T2S^del-P^ mutant (Fig. 2D). When the T2S^del-P^ and T2S^sil-P^ mutants served as the female strain in crosses to the *mat1-1-1* deletion mutant labeled with an H1-GFP, all outcrosses formed morphologically normal asci and ascospores with the expected 1:1 segregation of 8 ascospores with and without GFP signals in an ascus. These results indicate that the S-transcript of *FgTAD2* is expressed and functional in sexual dikaryotic tissues. Strand-specific RNA-seq analysis showed that mRNA editing activities were remarkably decreased relative to the wild type, and only an average of 36 and 265 A-to-I RNA editing sites were detected in 7-dpf perithecia of the T2S^del-P^ and T2S^sil-P^ mutants, respectively (Fig. 2E). As a control, an average of 327 and 388 A-to-I RNA editing sites were detected in the *mat1-1-1* and *mat1-2-1* deletion mutants, respectively, which produce small barren perithecia blocked in the development of ascogenous hyphae (*13*). The median editing levels of detected A-to-I sites in the T2S^del-P^ and T2S^sil-P^ mutants were also significantly reduced in comparison with the *mat* mutants. Therefore, the S-transcript of *FgTAD2* is critical for ascogenous hypha formation and A-to-I mRNA editing in *F. graminearum*.

Likewise, we removed the promoter region (−13 to −434 bp relative to the start codon) of the *FgTAD3* S-transcript at the native locus. The resulting T3S^del-P^ mutant was normal in vegetative growth and perithecium formation. Asci were normally formed in perithecia, but no ascospores were observed in asci (Fig. 2D). Both the number and median editing levels of A-to-I RNA editing sites were markedly decreased in the T3S^del-P^ mutant compared with the wild type (Fig. 2E), suggesting that the *FgTAD3* S-transcript is important for A-to-I mRNA editing. Since both FgTad2 and FgTad3 are essential for RNA editing, the relatively small effect caused by the deletion of the *FgTAD3* S-transcript suggests that the *FgTAD3* L-transcript is also expressed and functional in sexual dikaryotic tissues.

### The start codon of the *FgTAD2* S-transcript corresponds to the M^4^ codon of the L-transcript

To assay the protein products of *FgTAD2*, we generated transformants expressing a *FgTAD2*-FLAG fusion protein *in situ* and detected only one protein product in 7-dpf perithecia by western blot analysis (Fig. 2F). Likewise, only one protein product was detected in the transformants ectopically expressing a *FgTAD2*-FLAG construct under the control of the RP27 promoter (Fig. 2G). To determine which transcript the detected protein was expressed, we generated transformants ectopically expressing an allele with multiple synonymous mutations at the promoter region of the S-transcript. The abundance of detected proteins in the *FgTAD2*-S^sil-P^-FLAG transformant was remarkably decreased in perithecia compared with the transformant expressing the wild-type allele *FgTAD2*-FLAG (Fig. 2G), indicating that the detected protein in perithecia is mainly derived from the S-transcript.

The detected transcriptional initiation sites of the *FgTAD2* S-transcript were located before the fourth in-frame AUG codon (M^4^) of the L-transcript, suggesting that the M^4^ codon is most likely functional as the start codon of the S-transcript. To confirm the translational initiation of the detected proteins, we generated transformants ectopically expressing an M^4^-initiated *FgTAD2*-FLAG construct. The dominant protein product expressed by the M^4^-*FgTAD2*-FLAG transformant had the same size as the proteins detected in previous transformants (Fig. 2G), confirming that the *FgTAD2* S-transcript encodes a protein with a start codon corresponding to the M^4^ codon of the L-transcript. The M^1^-FgTad2 was hardly detected in the 7-dpf perithecia, possibly due to the low expression level of the L-transcript in the late stage of sexual development.

To verify its roles, we introduced an A-to-C mutation to the M^4^ codon *in situ*. The resulting *FgTAD2^M4L^*mutant was normal in vegetative growth but defective in ascospore formation (Fig. 2H). It formed fewer ascospores with morphological abnormalities in asci. In contrast, mutations of M^3^, M^6^, or M^7^ codon caused no obvious phenotypic changes (Fig. S1). Therefore, M^4^ is important for the function of the *FgTAD2* S-transcript. The defect caused by M^4^ mutations is possibly due to the low translation efficiency of the S-transcript when using the downstream M codon as the start codon.

### Alternative translational initiation generates two protein isoforms from the *FgTAD2* L-transcript

Unexpectedly, although only the L-transcript is expressed, two protein products were detected in hyphae (Fig. 2F-2G). 5’RACE confirmed that only the L-transcript was expressed in hyphae of the *in situ FgTAD2*-FLAG transformant. Given the protein size, the two protein products were most likely expressed initiated from the M^1^ and M^4^ codons, respectively. We then generated transformants ectopically expressing the *FgTAD2^M1L^*-FLAG and *FgTAD2^M4L^*-FLAG alleles by introducing an A-to-C mutation to the M^1^ and M^4^ codons, respectively. In the *FgTAD2^M1L^*-FLAG transformant, the M^1^-protein band completely disappeared, but beyond the M^4^-protein, two new bands potentially corresponding to the size of M^2^- and M^3^-proteins were detected (Fig. 2G). Only trace amounts of M^4^-proteins were detected in the *FgTAD2^M4L^*-FLAG transformant. The above results suggest that the L-transcript expresses two protein isoforms using the M^1^ and M^4^ codons as start codons, respectively.

M^1^-FgTad2 had a 112 aa N-terminal extension relative to M^4^-FgTad2. Both protein isoforms contained the entire CDA domain. To investigate the biological role of this N-terminal extension, we changed the first three M (ATG) codons into L (CTT) codons *in situ* simultaneously (Fig. 2H). The resulting *FgTAD2^M1,2,3L^* mutant expected to express only the M^4^-protein had no obvious phenotypic changes, indicating that the N-terminal extension of the M^1^-FgTad2 is dispensable for vegetative growth and sexual reproduction. However, Y2H assays revealed that M^4^-FgTad2 had a stronger interaction with FgTad3 compared with the M^1^-FgTad2 (Fig. 2I). Therefore, the interaction of the FgTad2-FgTad3 complex is negatively affected by the N-terminal extension of the M^1^-FgTad2.

### Sexual stage-specific post-translational modifications are important for the function of FgTad3

The two transcript isoforms of *FgTAD3* share the same ORF, and only the length of the 5’-untranslated regions (UTR) is different. Surprisingly, when a *FgTAD3*-GFP fusion protein was expressed under the control of the RP27 promoter in PH-1, two bands of smaller sizes were detected by western blot analysis in perithecia, instead of the expected 73 kDa band detected in hyphae (Fig. 2J). The size of these bands suggests that they are unlikely to be a result of the use of alternative in-frame AUG (M) codons, as observed for *FgTAD2*. It is possible that *FgTAD3* undergoes post-translational modifications specific to the sexual stage, which could explain this observation. Using tandem mass spectrometry (MS/MS) analysis, we identified four post-translational modification sites in FgTad3 that are uniquely present in perithecia compared to hyphae. These sites include one acetylated site (acK^198^), two non-canonical phosphorylated sites (pE^8^ and pE^241^), and one methylated site (meE^17^) (Fig. 2K and Fig. S2).

To verify the roles of these modifications, we generated FgTad3-hypomodified mutants by replacing each modified residue with either alanine (A) or arginine (R) to prevent potential modifications. While the *FgTAD3^E8A^* and *FgTAD3^E17A^* mutants had no detectable phenotypes (Fig. S2), the *FgTAD3^K198R^* and *FgTAD3^E241A^* mutants were normal in vegetative growth but defective in ascospore formation (Fig. 2L). Most of the asci in perithecia produced by the two mutants contained no ascospores at 7-dpf compared to the wild type. The E^241^ phosphorylation site is located within the CDA domain, while the K^198^ acetylation site is in the linker region that connects the FgTad3^CDA^ and FgTad3^N^ domains (Fig. 2M). These results suggest that the acK^198^ and pE^241^ modifications are important for the function of FgTad3.

We further generated *FgTAD3^K198Q^* and *FgTAD3^E241D^* mutants by replacing K^198^ and E^241^ with glutamine (Q) and aspartic acid (D), respectively, to mimic the acetylated and phosphorylated state of FgTad3. Both mutants had a slight defect in growth and formed a few perithecia on carrot agar (Fig. 2L). Only a small number of elongated asci but no ascospores were observed in the perithecia produced by the two mutants. These findings suggest that constitutive K^198^ acetylation and E^241^ phosphorylation harm growth and perithecium formation because the modifications are sexual stage-specific.

### Identifying a sexual stage-specific protein essential for A-to-I mRNA editing

As A-to-I mRNA editing also occurs during the sexual stage of *N. crassa*, it is expected that the putative factors that activate mRNA editing are specific to the sexual stage and conserved between *F. graminearum* and *N. crassa*. Using our previously published RNA-seq data (*11, 14*), we identified 34 orthologs that are expressed specifically during sexual development in both fungi (Table S1), which we named <u>s</u>exual stage-<u>s</u>pecific <u>c</u>onserved (SSC) genes. Out of these, eight genes have already been published and their reported phenotypes suggest that they are not associated with mRNA editing. For the remaining 26 SSC genes, we generated deletion mutants for each of them. As expected, all deletion mutants showed normal growth. However, deletion mutants of 6 SSC genes displayed defects in different stages of sexual development (Table S1). We focused on two deletion mutants that had defects arrested at earlier stages of perithecium development, namely *ssc23* (FG4G02710) and *ssc20* (FG3G23790). The *ssc23* mutant failed to produce perithecia while the *ssc20* mutant formed smaller perithecia with no visible ascogenous hyphae in them (Fig. 3A-3C). The complemented strains had normal phenotypes as the wild type. Through strand-specific RNA-seq analysis of 60-h sexual cultures, we identified an average of 152 A-to-I mRNA editing sites in the wild type and 26 in the *ssc23* mutant (Table S2). However, we did not detect any reliable A-to-I mRNA editing sites in the *ssc20* mutant, despite the *ssc23* mutant having more severe defects. Moreover, we did not find any reliable A-to-I mRNA editing sites in the 6-dpf perithecia of the *ssc20* mutant. These results suggest that *SSC20* is essential for A-to-I mRNA editing. As a result, we designated *SSC20* as *AME1* (activator of mRNA editing).

**Fig. 3.**
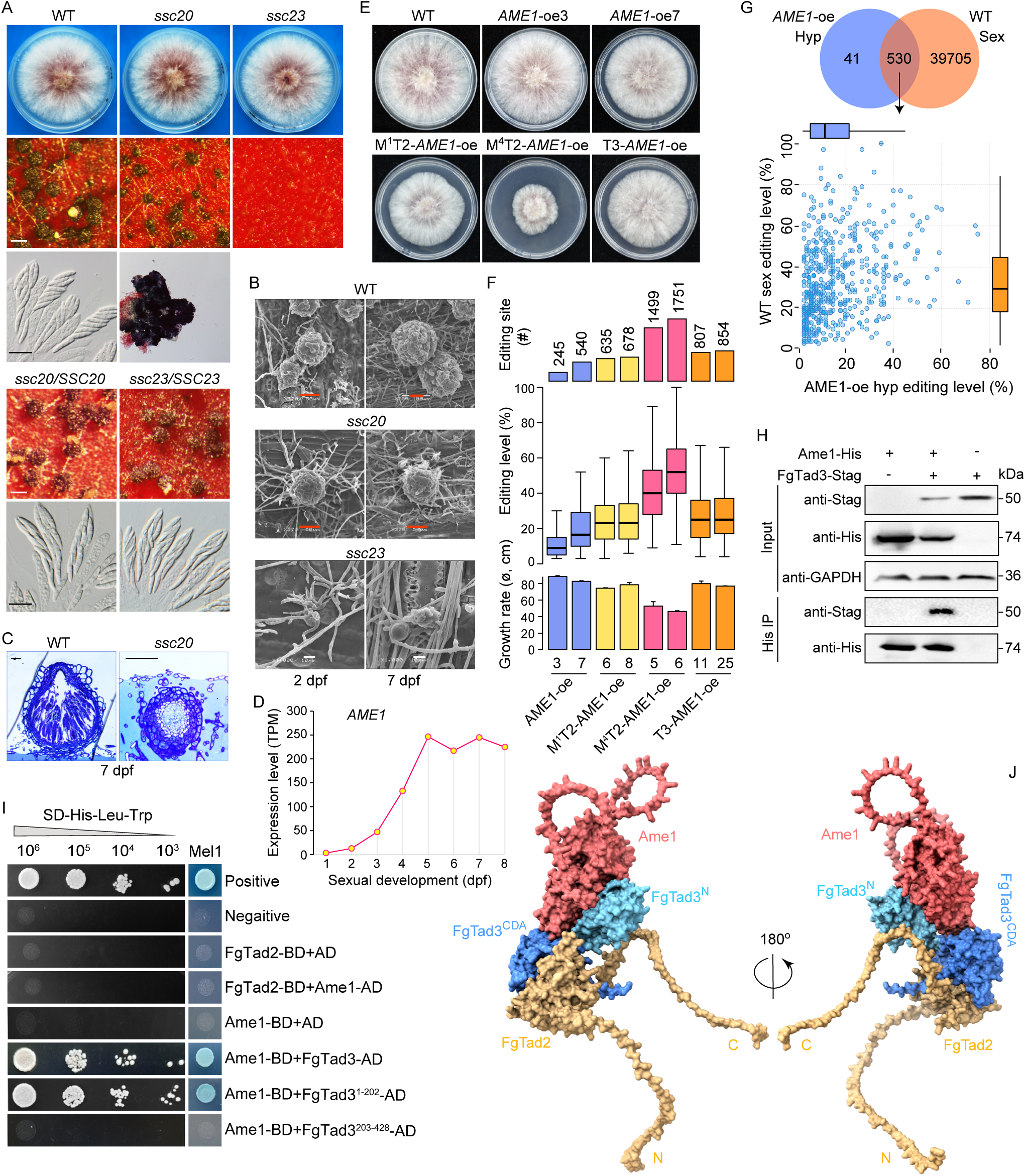
A sexual stage-specific protein Ame1 essential for sexual reproduction and A-to-I mRNA editing. (A) Colony morphology, perithecium formation, and asci/ascospores morphology of deletion mutants and complemented strains of *AME1/SSC20* and *SSC23*. White bar = 0.2 mm; black bar = 20 μm. (B) Scanning electron microscope of 2-dpf and 7-dpf perithecia produced on wheat straw. Red bar = 50 μm; white bar = 10 μm. (C) Semi-thin sections of 7-dpf perithecia. bar = 20 μm. (D) Expression levels of *AME1/SSC20* during sexual development from 1 to 8 dpf. (E) Colony morphology of transformants expressing *AME1* (*AME1*-oe), or co-expressing *AME1* with M^1^-*FgTAD2* or M^4^-*FgTAD2* (M^1^T2-*AME1*-oe or M^4^T2-*AME1*-oe), or with *FgTAD3* (T3-*AME1*-oe) under the control of the RP27 promoter. (F) The number of A-to-I mRNA editing sites detected in 24-h hyphae, the editing levels of common editing sites, and the colony growth rate of marked strains. For the bar graph, Mean and SD were calculated with data from three independent repeats (n = 3). (G) Comparison of editing levels of overlapped A-to-I mRNA editing sites detected in the hyphae of *AME1*-oe transformants and the perithecia of wild-type strains. (H) Co-IP assays for the interaction between Ame1 and FgTad3. Western blots of total proteins isolated from *E. coli* transformants expressing the Ame1-6×His and/or FgTad3-Stag (input) and proteins eluted from anti-His affinity beads (His IP) were detected with anti-His and anti-Stag antibodies. (I) Yeast two-hybrid assays for the interaction of Ame1 (BD-bait) with FgTad2 or FgTad3, as well as FgTad3 mutants (AD-prey). (J) Surface model of the ternary complex M^4^-FgTad2-FgTad3-Ame1 predicted with AlphaFold-Multimer.

### Expression of *AME1* in hyphae leads to widespread A-to-I mRNA editing

During sexual development, the expression of *AME1* was continuously upregulated from 1 dpf to 5 dpf, after which the expression level tended to stabilize (Fig. 3D), which is consistent with the high editing activity observed after 5 dpf (Fig. 1B). To further confirm the role of *AME1* in mRNA editing, we replaced its promoter with the RP27 promoter *in situ*. The resulting *AME1*-oe transformants showed phenotypic variations in vegetative growth, with some having wild-type phenotypes and others having slightly reduced growth rates (Fig. 3E). Interestingly, we detected plentiful A-to-I mRNA editing sites in the hyphae of both types of transformants, with 245 sites in a normal transformant and 540 sites in a defective transformant (Fig. 3F), demonstrating that Ame1 can activate mRNA editing.

The analysis revealed that more than 90% of the editing sites detected in hyphae were also edited in perithecia (Fig. 3G). These overlapping sites were highly edited in perithecia but had a low editing level in hyphae. We then overexpressed *FgTAD3* or the M^1^- and M^4^-initiated *FgTAD2* with the RP27 promoter at a targeted locus in the *AME1*-oe transformant. All resulting overexpressing transformants had growth defects, especially the M^4^T2-*AME1*-oe transformants whose growth rates were reduced by ∼50% (Fig. 3E). We detected 644 to 1760 A-to-I mRNA editing sites in these overexpressing transformants (Fig. 3F). The number of detected editing sites in vegetative growth remains much lower than that in the perithecia. The number and median editing level of editing sites detected in different transformants were correlated with the extent of their growth defects, suggesting that mRNA editing during vegetative growth is detrimental. The results also indicate that the M^4^-FgTad2 has higher editing activity, consistent with its stronger interaction with FgTad3.

### Interaction between Ame1 and the N-terminal domain of FgTad3

To investigate the relationship between Ame1 and FgTad2-FgTad3, we generated transformants ectopically expressing *FgTAD3*-FLAG, *AME1*-GFP, or both constructs and performed co-IP assays. While we detected FgTad3-FLAG and Ame1-GFP proteins in transformants where they were expressed individually, we hardly detected Ame1-GFP in the co-expressed transformant (Fig. S3). This suggests that expressing both proteins is detrimental to growth and is therefore repressed. We then transformed the *FgTAD3-*STAG, *AME1-*6×HIS, or both constructs into *Escherichia coli* and confirmed that Ame1 was associated with the FgTad2-FgTad3 complex *in vivo* using co-IP results (Fig. 3H). Based on Y2H assays, it was observed that Ame1 directly interacted with FgTad3 but not with FgTad2 (Fig. 3I). Furthermore, Ame1 directly interacted with the N-terminal domain (1-202 aa) of FgTad3, while no interaction was observed with the C-terminal domain (203-428 aa). Using AlphaFold-Multimer (*15*), we predicted heteromeric interfaces among M^4^-FgTad2, FgTad3, and Ame1, showing that the ternary complex contained two semi-independent domains. The first domain comprised mainly FgTad2 and the FgTad3 C-terminal domain, while the second domain comprised mainly Ame1 and the FgTad3 N-terminal domain (Fig. 3J). Contacting Ame1 to the N-terminal domain of FgTad3 changes the substrate recognition and specificity of the deaminase as the FgTad3 N-terminal domain is known for tRNA binding.

### Evolutionary origin of Ame1 and pivotal residues responsible for mRNA editing ability in Sordariomycetes

Ame1 is a protein of unknown function that contains a DUF726 domain belonging to the alpha/beta hydrolase superfamily (cl21494) (Fig. 4A). It also possesses the characteristic catalytic triad (S-D-H) active site found in other members of this superfamily. We constructed mutants in PH-1 to investigate whether the function of Ame1 on mRNA editing depends on the catalytic triad. Mutants with S^531^ and H^629^ mutations (*AME1^S531A^* and *AME1 ^S531A,H629A^*) showed normal sexual development, while the *AME1^D589A^* mutant formed small perithecia similar to the *ame1/ssc20* deletion mutant (Fig. 4B), indicating that the catalytic activity is dispensable for Ame1’s function in mRNA editing. Y2H assays did not detect the interaction between Ame1^D589A^ and FgTad3 (Fig. 4C), suggesting that D^589^ is important for the interaction with FgTad3.

**Fig. 4.**
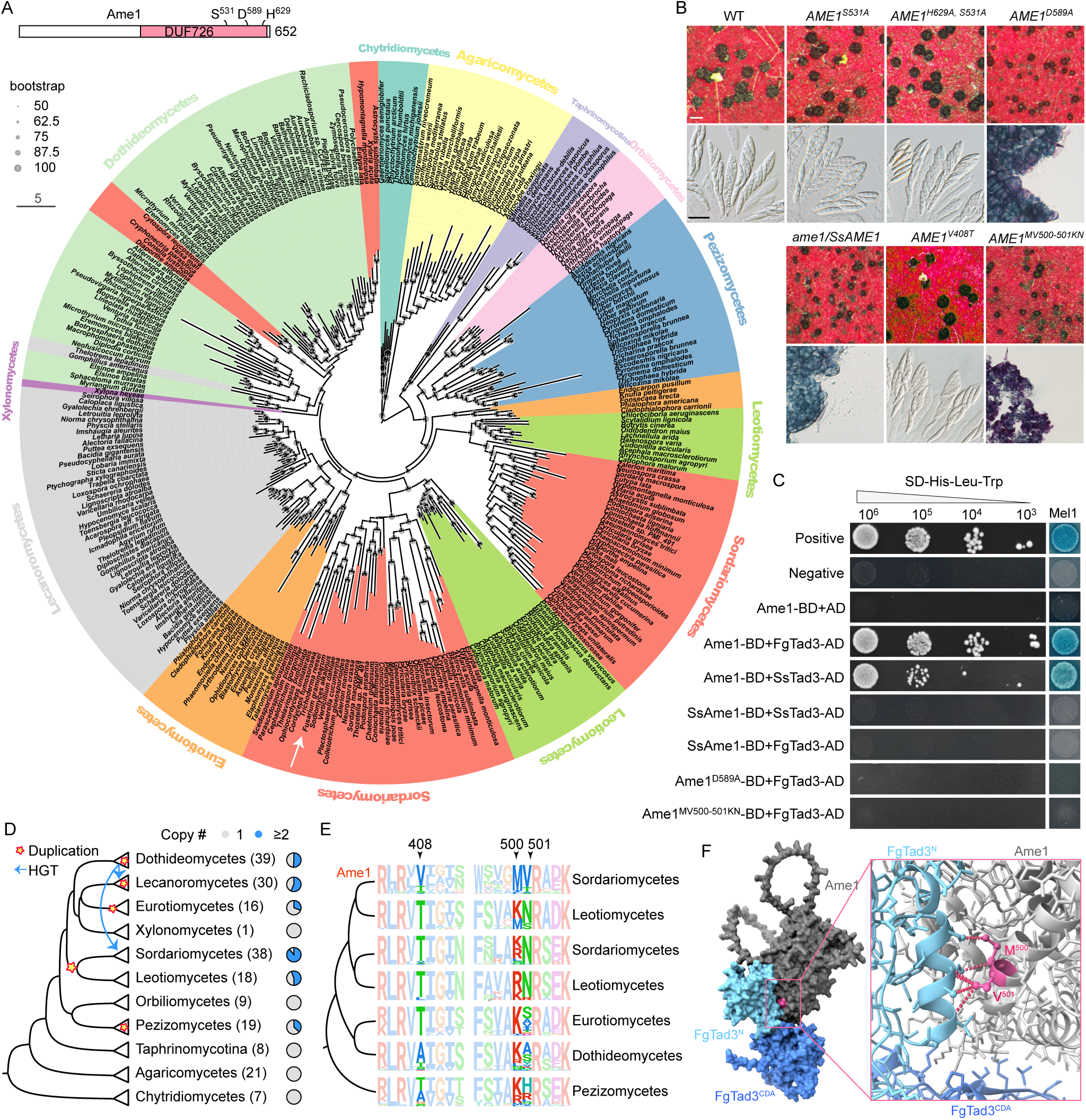
Evolutionary origin of Ame1 and key residues responsible for mRNA editing ability in Sordariomycetes. (A) The protein domain of Ame1 and a Maximum Likelihood phylogenetic tree of Ame1 orthologs from different fungal lineages. The putative catalytic triad (S-D-H) active sites are indicated. The white arrow indicates the position of *AME1* in the tree. (B) Perithecium formation and asci/ascospores morphology of the marked *AME1* mutants. White bar = 0.2 mm; black bar = 20 μm. (C) Yeast two-hybrid assays for the interaction among Ame1, Ame1 mutants, FgTad3, and their orthologs SsAme1 and SsTad3 from *Sclerotinia sclerotiorum*. (D) The dendrogram of different fungal taxa. The number of species in each taxon examined is indicated in the bracket. Each pie chart shows the proportion of fungal species with 1 or ≥2 copies of *AME1* orthologs derived from gene duplication and/or horizontal gene transfer events. The putative origin of gene duplication and horizontal gene transfer events are indicated in the dendrogram. (E) Sequence logos illustrate the conservation of amino acid residues within different fungal taxa at three putative sites responsible for the RNA editing ability of Ame1. (F) Direct contact between the MV^500-501^ residues of Ame1 and the interface residues of FgTad3 shown in the protein model.

Ame1 orthologs are widely distributed in ascomycetes but have been lost in Saccharomycotina (Fig. 4A). Orthologs have also been detected in Agaricomycetes and Chytridiomycetes. Multiple independent gene duplication and horizontal gene transfer events have occurred in different lineages in filamentous ascomycetes, with an ancestral duplication event giving rise to two copies at least in the last common ancestor of Sordariomycetes and Leotiomycetes (Fig. 4D). The branch length of Ame1 in Sordariomycetes was notably longer than that in Leotiomycetes (Fig. 4A), indicating accelerated evolution of Ame1 in Sordariomycetes. As A-to-I mRNA editing is a common feature in Sordariomycetes but has not been found in Leotiomycetes yet, we hypothesized that the function of Ame1 in mRNA editing has evolved in Sordariomycetes. To test this hypothesis, we replaced the *AME1* ORF *in situ* in PH-1 with the ORF of the *AME1* ortholog *SsAME1* from *Sclerotinia sclerotiorum*, a species in Leotiomycetes. The resulting *ame1/SsAME1* transformants had similar defects as the *ame1* deletion mutant (Fig. 4B), indicating that the mRNA editing function of *AME1* cannot be replaced by *SsAME1*. The Y2H assay showed that SsAme1 did not interact with both FgTad3 and SsTad3, but Ame1 did interact with SsTad3 (Fig. 4C). This indicates that the capability of Ame1 to interact with Tad3 evolved in Sordariomycetes.

We identified three amino acid sites that may potentially contribute to the functional innovation of Ame1 in Sordariomycetes (Fig. 4E). We created mutants for these sites *in situ* in PH-1 by replacing the conserved residues in Sordariomycetes with those in Leotiomycetes. The *AME1^V408T^* mutant displayed normal phenotypes, while the *AME1^MV500-501KN^* mutant exhibited similar defects to the *ame1* deletion mutant (Fig. 4B), indicating that MV^500-501^ is essential for Ame1’s function in mRNA editing. Based on the protein structural model, the MV^500-501^ residues are located at the interaction interface between Ame1 and the N-terminal domain of FgTad3 and directly contact the interface residues of FgTad3 (Fig. 4F). Y2H assays confirmed that MV^500-501^ is crucial for the interaction of Ame1 with FgTad3 (Fig. 4C). Because the ancestors of Ame1 had KN residues at this position, the replacement of KN with MV in Sordariomycetes was a critical step in the evolution of Ame1’s ability to interact with Tad3 and enable mRNA editing.

### Amino acid residues in FgTad3 uniquely important for mRNA editing

To identify the amino acid residues of FgTad3 that are uniquely important for mRNA editing, we used the repeat-induced point mutation (RIP) mechanism to generate mutations in FgTad3. This mechanism generates C-to-T mutations randomly in repetitive sequences premeiotically (*16*). The coding region of *FgTAD3* without the start codon was inserted in a reverse orientation before the promoter region of *FgTAD3 in situ*, aiming to generate tandemly repeated *FgTAD3* without affecting the transcription of the native gene (Fig. 5A). The resulting TR-*FgTAD3* transformants exhibited normal vegetative growth. However, during sexual reproduction, a few asci with abnormal ascospores were observed (Fig. 5B), indicating that RIP mutations had occurred in *FgTAD3*. We isolated ascospores with abnormal morphology at 10 dpf and obtained a total of 350 ascospore progeny. Among these, 33 showed normal growth but exhibited varying degrees of defects in sexual development (Table S3). As the inserted *FgTAD3* segment is not transcribed, we amplified and sequenced the native *FgTAD3* gene. All ascospore progeny had at least one RIP mutation resulting in amino acid changes in FgTad3. A total of 52 non-synonymous RIP mutation sites were identified, with most of them located in the N-terminal part of FgTad3 (Table S3).

**Fig. 5.**
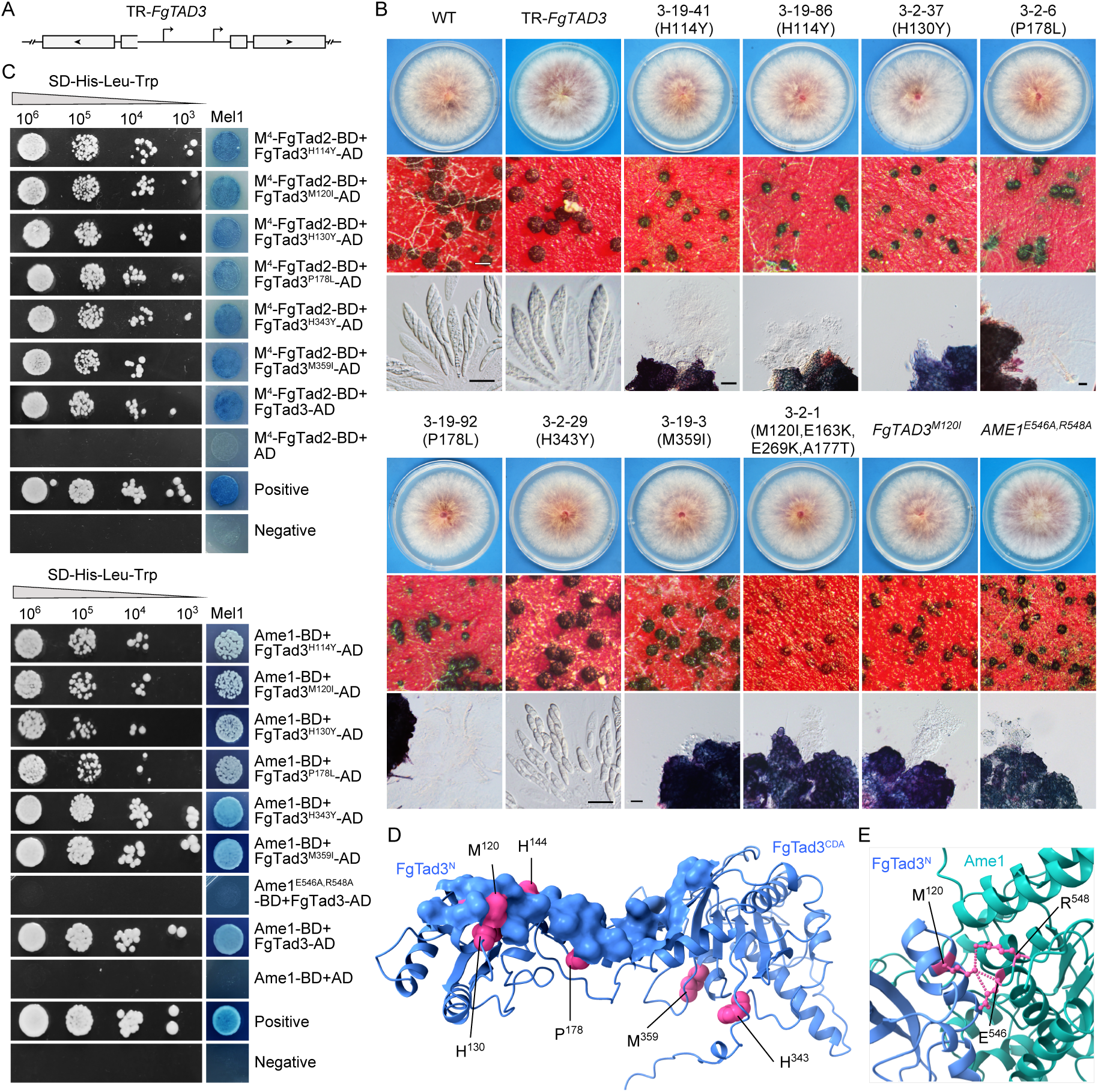
Amino acid residues in FgTad3 uniquely important for sexual reproduction and interaction with Ame1. (A) Schematic representation of tandemly repeated *FgTAD3* construct in TR-*FgTAD3* strain used for RIP mutations of *FgTAD3*. Arrows indicate transcriptional initiation sites and boxes indicate CDS regions. (B) Colony morphology, perithecium formation, and asci/ascospores morphology of the TR-*FgTAD3* strain, its ascospore progeny with RIP mutations, and the marked mutants of *FgTAD3* and *AME1*. White bar = 0.2 mm; black bar = 20 μm. (C) Yeast two-hybrid assays for the interaction of the marked mutant alleles. (D) Location of mutated residues in proximity to contacting surface of FgTad3 with Ame1. (E) Direct contact between Ame1 E^546^R^548^ residues and FgTad3 M^120^ residue shown in the protein model.

Among the ascospore progeny, seven showed normal growth but severe defects in sexual development (Fig. 5B), each containing a single non-synonymous RIP mutation in *FgTAD3*. Therefore, the observed defects can be attributed to the specific mutational effects. Out of these progeny, progeny 3-2-29 (H343Y) displayed defects only in ascospore formation. The other six progeny produced small perithecia, but no asci were observed in the perithecia formed by progeny 3-19-41 and 3-19-86 (both carrying the H114Y mutation), 3-2-37 (H130Y), and 3-19-3 (M359I). Only a few elongated asci without ascospores were observed in the perithecia formed by progeny 3-19-92 and 3-2-6 (both carrying the P178L mutation). Notably, among the progeny analyzed, the M120I mutation was the most frequent, observed in 9 progeny, which also carried additional mutations. To investigate the specific effect of the M120I mutation, we generated the mutation *in situ* in PH-1. The resulting *FgTAD3^M120I^*mutant exhibited normal growth but formed small perithecia without ascogenous hyphae (Fig. 5B). Only 40 A-to-I RNA editing sites were identified in 7-dpf perithecia of *FgTAD3^M120I^* mutants (Table S2), confirming the critical role of M^120^ in mRNA editing. Interestingly, 6 of the progeny with the M120I mutation exhibited small perithecia without ascogenous hyphae, similar to what was observed in *FgTAD3^M120I^* mutants (Fig. 5B). However, a few ascogenous hyphae could be visible inside the slightly small perithecia formed by progeny 3-2-50, 3-19-94, and 3-19-50, which all contain an additional V82I mutation (Table S3). These results suggest that the V82I mutation may have a compensatory effect on the M120I mutation.

Y2H assays showed that the mutant alleles FgTad3^H114Y^, FgTad3^M120I^, FgTad3^H130Y^, and FgTad3^P178L^ had a normal interaction with M^4^-FgTad2 but a decreased interaction with Ame1(Fig. 5C), suggesting that these sites in FgTad3 are important for the interaction with Ame1. Consistently, residues H^144^, M^120^, H^130^, and P^178^ are located within or close to the interacting surface of FgTad3 and Ame1 (Fig. 5C). On the other hand, the interaction with both Ame1 and FgTad2 did not show any obvious changes for FgTad3^H343Y^ and FgTad3^M359I^ (Fig. 5C). The H343Y and M359I mutations are located in the CDA domain of FgTad3 and may affect the conformation of the enzyme that is required for accommodating the stem-loop structures of mRNA.

We identified two residues, E^546^ and R^548^, in Ame1 that were directly in contact with M^120^ of FgTad3 (Fig. 5E). To investigate the significance of these residues, we introduced E546A and R548A mutations *in situ* in PH-1. The *AME1^E546A,R548A^* mutant exhibited normal growth but formed small perithecia without ascogenous hyphae (Fig. 5B). Y2H assays confirmed that no interaction was observed between Ame1^E546A,R548A^ and FgTad3, thus highlighting the critical role of E^546^R^548^ in the interaction between Ame1 and FgTad3.

### The FgTad2-FgTad3-Ame1 complex exhibits A-to-I mRNA editing activities in yeasts, bacteria, and human cell lines

To investigate the A-to-I mRNA editing activities of the FgTad2-FgTad3-Ame1 complex in heterologous systems, we co-expressed M^4^-*FgTAD2*, *FgTAD3*, and *AME1* in the *S. cerevisiae* yeast strain, INVSc1 under the control of the *GAL1* promoter. As controls, we also co-expressed M^4^-*FgTAD2* and *FgTAD3* or expressed only *AME1*, as well as co-expressed *FgTAD3*, *AME1*, and a mutant allele M^4^-*FgTAD2* with the active site E^121^ mutated into A (M^4^-*FgTAD2^E121A^*). To investigate the A-to-I mRNA editing activities of the FgTad2-FgTad3-Ame1 complex in heterologous systems, we expressed M^4^-*FgTAD2*, *FgTAD3*, and *AME1* together under the control of the *GAL1* promoter in the *S. cerevisiae* yeast strain, INVSc1. As controls, we also expressed M^4^-*FgTAD2* and *FgTAD3* together, expressed only *AME1*, and co-expressed *FgTAD3*, *AME1*, and a mutant allele M^4^-*FgTAD2* with the active site E^121^ mutated into A (M^4^-*FgTAD2^E121A^*). The self-editing sites in the transcripts of *FgTAD3* and *AME1* were used as targets to assess the editing activity. By analyzing the RT-PCR products amplified from the total RNA isolated from the yeast transformant co-expressing M^4^-*FgTAD2*, *FgTAD3*, and *AME1*, we observed two peaks (A and G) at the target editing sites in Sanger sequencing traces (Fig. 6A). In contrast, only one A peak was detected at the target sites in the yeast transformant co-expressing M^4^-*FgTAD2^E121A^*, *FgTAD3*, and *AME1*, co-expressing M^4^-*FgTAD2* and *FgTAD3* or expressing only *AME1*. These results indicate that the FgTad2-FgTad3-Ame1 complex can perform A-to-I mRNA editing at the target sites in yeast, and the E^121^ in *FgTAD2* is crucial for editing. We also tested the editing activity of this complex in human HEK 293T cell lines. A-to-I mRNA editing was detected at the target sites of *FgTAD3* and *AME1* in the HEK 293T cells, with high editing activities. In contrast, editing was not detected at the target sites in the cells co-expressing M^4^-*FgTAD2* and *FgTAD3* or expressing only *AME1*. These results suggest that the FgTad2-FgTad3-Ame1 complex has universal activity in eukaryotic cells. Furthermore, we tested the editing activity of this complex in the *E. coli* BL21 strain using codon-optimized sequences. A-to-I mRNA editing was detected at a site in the *FgTAD3* transcript in the BL21 transformant co-expressing all three genes but not in the transformant co-expressing only M^4^-*FgTAD2* and *FgTAD3*. This suggests that the FgTad2-FgTad3-Ame1 complex also has mRNA editing activity in prokaryotic cells.

**Fig. 6.**
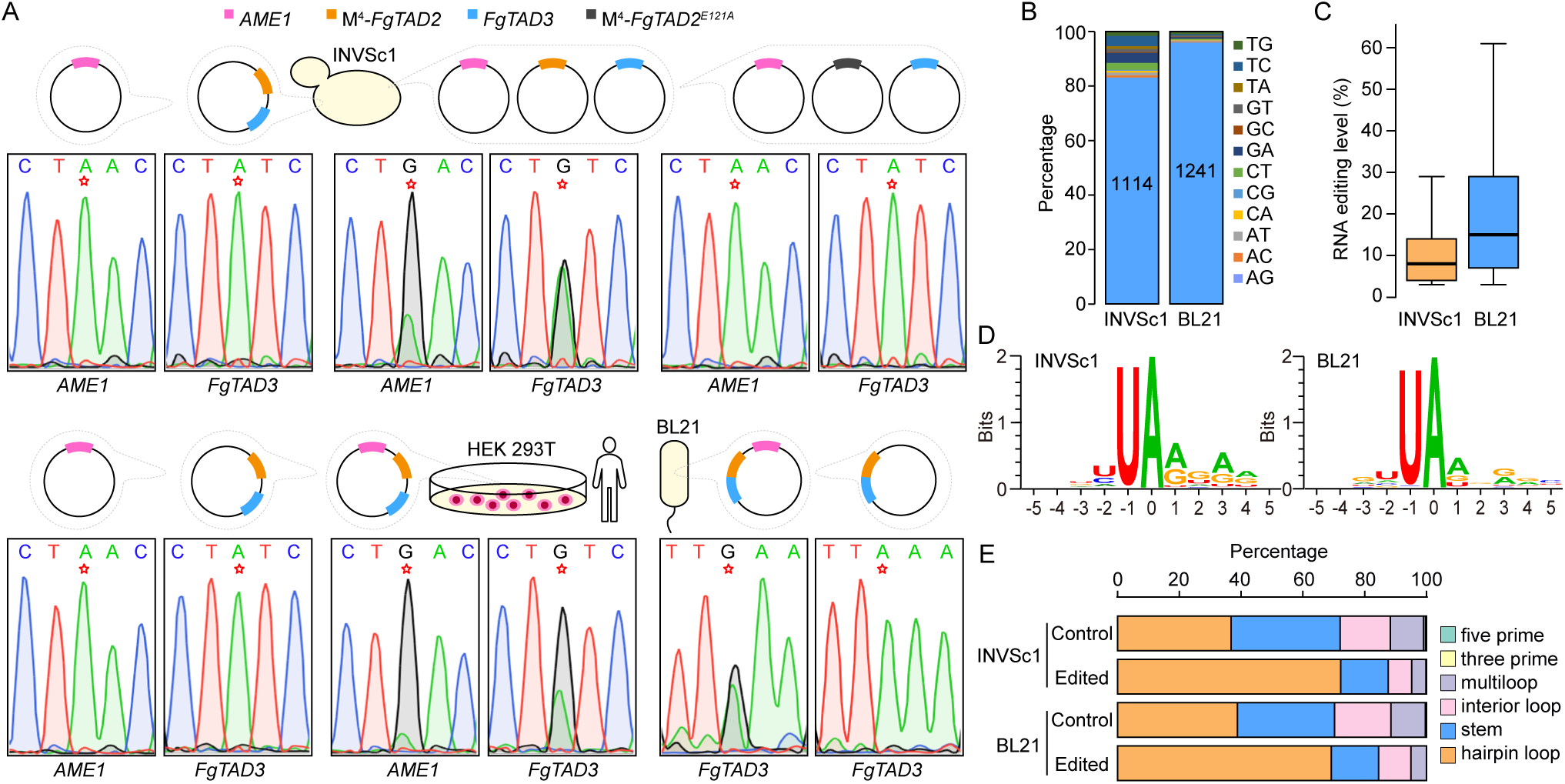
A-to-I mRNA editing activities of the FgTad2-FgTad3-Ame1 complex in heterologous systems. (A) Schematic representation of the expression system used for transformation of the yeast INvSC1 strain, the *E. coli* BL21 strain, and the human HEK 293T cell line. Self-editing of the transcripts of *AME1* and *FgTAD3* was detected by Sanger sequencing of the RT-PCR products. The target editing sites are marked by a red star. When editing occurs, two peaks (A and G) at the editing sites are observed in the sequencing traces. (B and C) The number and editing level (%) of detected A-to-I RNA editing sites in the yeast INvSC1 strain and *E. coli* BL21 strain expressing the FgTad2-FgTad3-Ame1 complex. (D) WebLogo showing base preferences in the flanking sequences of the editing sites. (E) Stacked columns showing fractions of different types of RNA secondary structure elements predicted based on 30-nt upstream and 30-nt downstream sequences of the edited A sites. For control, an equal number of A sites with similar nucleotide preference at the - 2 to +4 positions were randomly selected from transcript sequences.

**Fig. 7.**
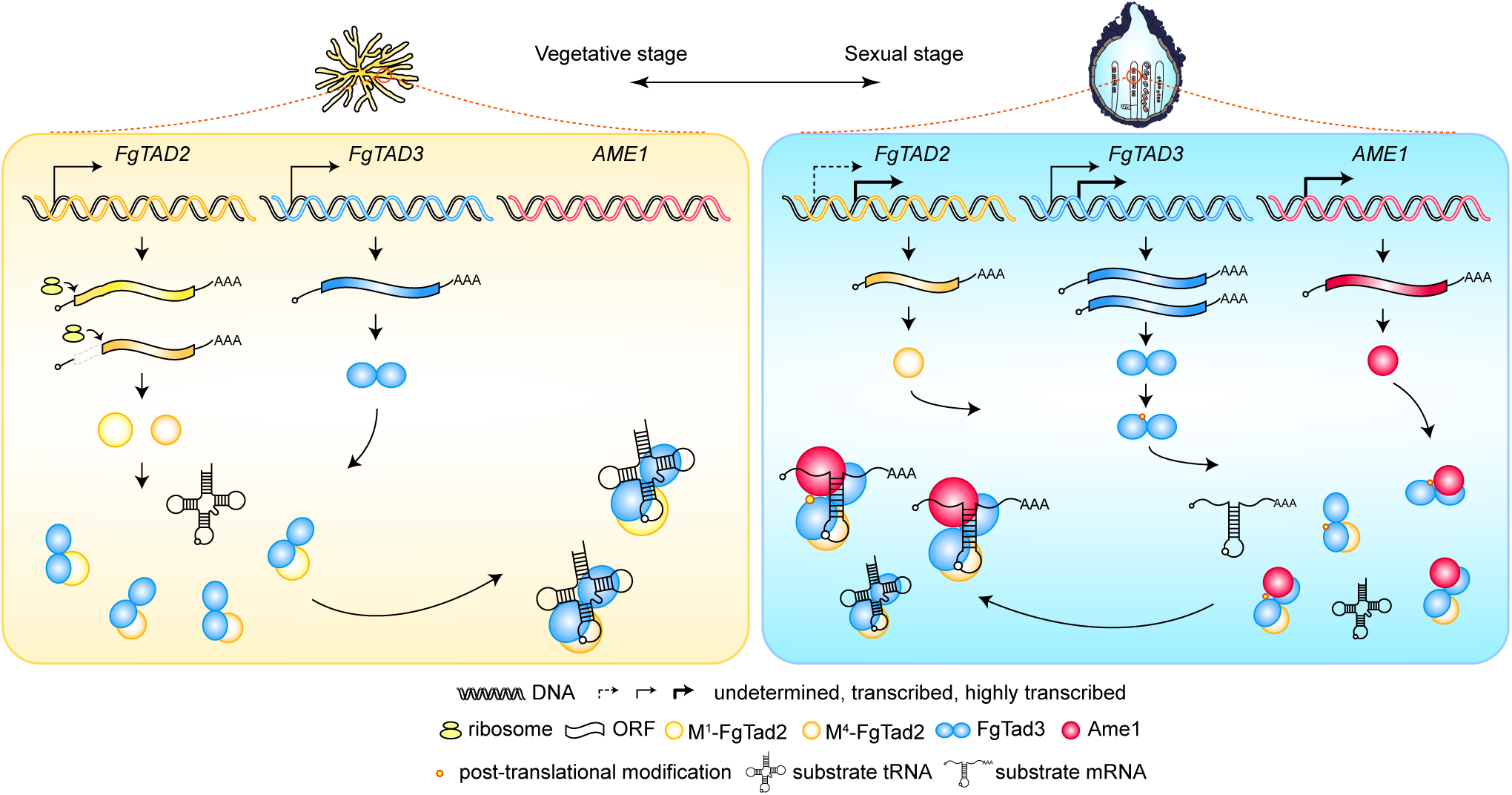
Proposed models for components, expression, and regulation of A-to-I RNA editing machinery in *F. graminearum*. *FgTAD2* encodes proteins that contain a CDA domain, while *FgTAD3* encodes proteins that contain both an N-terminal RNA-binding domain and a C-terminal CDA domain. FgTad2 interacts with the CDA domain of FgTad3 to form the catalytic domain responsible for A-to-I editing of both tRNAs and mRNAs. During vegetative stages, *FgTAD2* expresses two protein isoforms: M^1^-FgTad2 and M^4^-FgTad2, via alternative translational initiation. Compared to M^1^-FgTad2, M^4^-FgTad2 has a stronger interaction with FgTad3. In sexual stages, both *FgTAD2* and *FgTAD3* express a short transcript isoform via alternative transcriptional initiation in sexual dikaryotic tissues. The long transcript isoform of *FgTAD3* is also expressed, while that of *FgTAD2* remains to be determined. The short transcript isoform of *FgTAD2* encodes proteins identical to M^4^-FgTad2, while both transcript isoforms of *FgTAD3* encode the same proteins. A gene named *AME1* (activator of mRNA editing) is induced to expression in sexual stages, which encodes proteins with a DUF726 domain. Ame1 interacts with the N-terminal domain of FgTad3 to mediate the recognition of mRNA substrates. The FgTad2-FgTad3-Ame1 complex performs A-to-I mRNA editing. FgTad3 also undergoes post-translational modification that is important for mRNA editing.

To evaluate the editing of endogenous transcripts in the INVSc1 and BL21 strains, we conducted DNA-seq and strand-specific RNA-seq analyses. We identified a total of 1114 and 1241 A-to-I mRNA editing sites in the endogenous mRNAs in the INVSc1 and BL21 transformants co-expressing all three genes (Fig. 6B), respectively. Notably, the editing levels of the detected editing sites in the BL21 transformants were significantly higher than those detected in the INVSc1 transformants (Fig. 6C). We observed similar nucleotide preference surrounding the editing sites and enrichment of editing sites in hairpin loop structures for the editing sites detected in both INVSc1 and BL21 transformants (Fig. 6D-6E), as previously reported in *F. graminearum* (*12*). Therefore, the FgTad2-FgTad3-Ame1 complex exhibits robust A-to-I mRNA editing activities in heterologous systems.

## Discussion

A-to-I RNA editing systems have recently gained attention for their ability to recode genetic information at the mRNA level. This allows for the correction of pathogenic mutations and the reprogramming of genetic information, which is particularly relevant as C:G to T:A genetic mutations account for approximately half of all known pathogenic point mutations in humans (*17, 18*). Site-directed RNA editing systems have been developed using ADARs to target disease-causing point mutations (*19, 20*). However, the eukaryotic tRNA-specific heterodimeric deaminase Tad2-Tad3 requires the complete tertiary structure of cognate tRNAs to perform its deamination reaction and is not known to have mRNA editing capacity (*21, 22*). In our study, we have shown that the single Ame1 protein enables FgTad2-FgTad3 to perform transcriptome-wide A-to-I mRNA editing through interaction with the N-terminal domain of FgTad3. This interaction expands the substrate recognition and specificity of the tRNA-specific deaminase from tRNA to include mRNA. While other regulatory factors may contribute to the efficiency of editing, our findings suggest that the FgTad2-FgTad3-Ame1 ternary complex is sufficient for efficient A-to-I editing of mRNA and may represent a minimal mRNA editing complex. While it is currently unknown if this complex retains the ability to edit tRNA, we have identified specific amino acid mutations in FgTad3 that impact mRNA editing. This helps to elucidate the differences in the mechanisms of tRNA-substrate and mRNA-substrate engagement by this complex. Previous reports have shown that altering specific residues through directed evolution can modify the substrate recognition and specificity of a deaminase (*23–25*). Our work provides a new perspective on how to control the substrate recognition and specificity of editing enzymes by utilizing their interaction with other partners. The ability of the fungal editing machinery to efficiently edit mRNA in different heterologous systems, including human cell lines, makes it a promising tool in therapy and agriculture. The fungal editing system and ADARs, which have different substrate preferences, can complement each other in RNA editing applications. Additionally, the preference of the fungal editing machinery for the UAG triplet (*8*) makes it especially useful for correcting nonsense mutations, which are responsible for a significant proportion of human inherited diseases (*26*).

Our study has shown that Ame1 is specific to the sexual stage and is directly responsible for the sexual stage-specific activity of A-to-I mRNA editing in fungi. The *ame1* deletion mutant produced small perithecia with no visible ascogenous hyphae, which is similar to the uneditable mutant of a single editing site in *PSC58*. This suggests that Ame1’s primary function is likely in mRNA editing, although other functions cannot be ruled out. We found that while A-to-I mRNA editing is essential for sexual development, activating it during vegetative growth can have harmful effects. The edited versions of mRNA seem to be beneficial in sexual stages but harmful in vegetative stages. This suggests that genomic mutations that advance sexual reproduction without distressing growth are unlikely due to the genetic trade-offs involved. The formation of the FgTad2-FgTad3-Ame1 complex specific to the sexual stage might provide an adaptive advantage by enabling a balance between reproduction and survival without compromising either. *F. graminearum* is a major cause of Fusarium head blight (FHB), one of the most devastating diseases on wheat and barley worldwide. Ascospores discharged from perithecia are the primary inoculum of FHB (*27*). Targeting the fungi-specific RNA editing machinery will be a promising approach to controlling fungal plant pathogens that rely on ascospores for infection (*28, 29*).

In addition to the sexual stage-specific interaction with Ame1, we have discovered that FgTad2-FgTad3 is regulated by alternative promoters, alternative translation initiation, and post-translational modifications. This indicates that the expression and activity of FgTad2-FgTad3 are tightly controlled within the cell. Notably, the M^4^-FgTad2 protein encoded by the sexual stage-specific S-transcript exhibits higher editing activity due to a stronger interaction with FgTad3 compared to the M^1^-FgTad2 protein encoded by the L-transcript. Surprisingly, the M^4^-FgTad2 isoform is also expressed by the L-transcript during growth through alternative translation initiation. The initiation of translation on eukaryotic mRNAs typically follows a scanning mechanism, where the first AUG encountered is favored as the start codon. However, when the first AUG codon is not in an optimal context, inefficient recognition of this codon can result in the preinitiation complex continuing to scan and initiating translation at a downstream AUG codon (*30*). Therefore, the M^1^ codon of the *FgTAD2* L-transcript may not be optimized relative to the M^4^ codon. Furthermore, the M^1^-FgTad2 isoform has an unusually long N-terminal part compared to orthologs in yeast, plants, and animals, suggesting that M^4^-FgTad2 represents the ancestral state. It is known that long 5’UTR transcripts tend to have lower translation efficiency (*31*). Because randomly generated mRNA editing activity is not necessarily specific to sexual reproduction at first, we speculate that the L-transcript of *FgTAD2* and *FgTAD3* may have evolved as a derived trait to mitigate the negative effects of mRNA editing during vegetative stages. However, further evidence is needed to support this hypothesis, which would provide insights into the regulatory evolution of A-to-I mRNA editing machinery in fungi.

Our study has shown that the role of Ame1 in mRNA editing has evolved in the last common ancestor of Sordariomycetes due to accelerated evolution after gene duplication, despite the widespread distribution of Ame1 orthologs in different fungal lineages. We have identified the key residues in Ame1 responsible for the innovation of mRNA editing in Sordariomycetes. Interestingly, sexual stage-specific A-to-I mRNA editing has also been identified in *Pyronema confluens* (*omphalodes*) (*32*), an early-diverging filamentous ascomycete belonging to Pezizomycetes. Our results suggest that A-to-I mRNA editing in the two fungal lineages has independent origins. Notably, *P. confluens* and its closely related lineages in Pezizomycetes also contain an additional duplicated copy of Ame1 orthologs, which raises the possibility that A-to-I mRNA editing in Pezizomycetes has also emerged due to the duplication of Ame1 orthologs. This work provides a strong motivation for future investigations of A-to-I mRNA editing in fungal lineages with duplication and horizontal gene transfer events in Ame1 orthologs.

## Acknowledgments

We thank Dr. Guanghui Wang, Ping Xiang, and Zhe Tang for their invaluable laboratory assistance. We thank Professor Junfeng Liu from China Agricultural University for generously providing us with the prokaryotic expression vector pRSFDuet-1. We also thank lab technicians Qiong Zhang and Xiaona Zhou from the State Key Laboratory for Crop Stress Resistance and High-Efficiency Production for their assistance with the MS/MS analysis. This work was supported by grants from the National Key R&D Program of China (2022YFD1400102) and the National Natural Science Foundation of China (no. 32170200).

## Author contributions

H.L. conceived and designed the experiments. C.F. performed experiments related to the identification of Ame1 and RIP mutations. K.X. performed experiments on FgTad2-overexpression and RIP-seq. K.X., J.Z., and C.F. performed experiments on transcriptional, translational, and post-translational regulation of FgTad2-FgTad3. Y.D. performed evolutionary analysis and functional site identification of Ame1, as well as bioinformatics analysis. Y.D., C.F., and R.Z. performed experiments related to assaying editing activities in heterologous systems. C.F., Y.D., K.X., X.X., Q.X., and Y.Z. performed the Y2H and co-IP experiments. H.L., J-R.X, C.F., Y.D., K.X., W.H., Q.W., C.J., X.W. and Z.K. analysed the data. H.L., C.F., K.X., Y.D, and J-R.X wrote the paper.

## Methods

### Strains and cultural conditions

The *F. graminearum* wild-type strain PH-1 (*33*) and its derived mutants and transformants were routinely cultured on potato dextrose agar (PDA) plates (20% potato, 2% dextrose, and 1.5% agar) at 25°C. Protoplast preparation and PEG-mediated transformation were performed as described previously (*34*). For transformant selection, hygromycin B (H005, MDbio, China) and geneticin (Sigma-Aldrich, St. Louis, MO) were added to the final concentration of 300 μg/ml and 200 μg/ml, respectively, in Top agar medium (0.3% yeast extract, 0.3% casamino acids, 20% sucrose, and 1.5% agar). The growth rate and colony morphology were assayed on PDA plates for three days. Vegetative hyphae used for DNA and RNA isolation were harvested from 24-hour liquid YEPD cultures (0.3% yeast extract, 1% peptone, and 2% dextrose). Conidia were harvested from 5-day-old liquid carboxymethyl cellulose (CMC) cultures (1.5% carboxymethylcellulose, 0.1% NH_4_NO_3_, 0.1% KH_2_PO_4_, 0.05% MgSO_4_·7H_2_O, and 0.1% yeast extract). For self-fertilization of sexual reproduction, aerial hyphae on 6-day-old carrot agar plates were pressed down with 500 μL of sterile 0.1% Tween-20 and subsequently cultured under black light at 25°C. For outcrosses, the mutant strains grown on carrot agar plates were fertilized with 1 mL of conidial suspension (1×10^6^ conidia/mL) derived from the *mat1-1-1* deletion mutant labeled with an H1-GFP. Perithecium formation and cirrhi production were examined using an Olympus SZX16 stereoscope. Ascogenous hyphae, asci, and ascospores were observed with an Olympus BX-51 microscope in squash mounts of perithecia 7 days post-fertilization (dpf). For scanning electron microscopy, perithecia produced on wheat straw were coated with gold-palladium and examined with a JEOL 6360 scanning electron microscope (Jeol).

### Gene deletion and complementation

Gene deletion was performed using the split-marker approach (*35*). Fragments of approximately 1.0 kb upstream and downstream of the target genes were amplified and connected to the N- and C-terminal regions of either a hygromycin-resistance cassette or a recyclable marker module (*12*) that confers resistance to hygromycin and sensitivity to the nucleoside analog 5-fluoro-2’-deoxyuridine (Floxuridine) through overlapping PCR. After transforming PH-1 protoplasts, hygromycin-resistant transformants were screened and confirmed by PCR assays. At least two independent deletion mutants were obtained for each gene. For gene complementation, the full-length genes, including their native promoter, were amplified and cloned into a pFL2 vector using the yeast gap-repair approach (*36*). The complementation construct was confirmed by DNA sequencing and then transformed into the protoplasts of the corresponding deletion mutants. Transformants containing the complementation constructs were screened with geneticin and confirmed by PCR assays. Table S4 lists all the strains used in this study, while Table S5 lists the primers used.

### Overexpression of FgTAD2 and FgTAD3 in F. graminearum

To overexpress the *FgTAD2* and *FgTAD3* genes in a target locus, the *FG1G36140* deletion mutant was used, which was generated by the recyclable marker module and had indistinguishable phenotypes from the wild type. The coding regions of *FgTAD2* or *FgTAD3* were PCR amplified and connected to the constitutive RP27 promoter from the pFL2 vector through overlapping PCR. Fragments of approximately 1.0 kb upstream and downstream of the *FG1G36140* gene were PCR amplified and connected to the P_RP27_-*FgTAD2* or P_RP27_-*FgTAD3* fragments through overlapping PCR. The PCR products were confirmed by sequencing analysis and then transformed into protoplasts of the *FG1G36140* deletion mutant. Transformants resistant to 25 μg/mL Floxuridine (HY-B0097, MCE, USA) were screened, as described (*12*). To overexpress *FgTAD3* or *FgTAD2* initiated by M^1^ or M^4^ codons in the *FG1G36140* locus of *AME1*-oe transformants, two DNA fragments were generated using overlapping PCR. One DNA fragment contained the upstream homologous fragment of *FG1G36140*, along with either the P_RP27_-*FgTAD3*, P_RP27_-M^1^-*FgTAD2*, or P_RP27_-M^4^-*FgTAD2* fragment, and the N-terminal region of the geneticin-resistance cassette. The other DNA fragment contained the C-terminal region of the geneticin-resistance cassette and the downstream homologous fragment of *FG1G36140*. These two DNA fragments were then co-transformed into the protoplasts of the *AME1*-oe transformants. Transformants resistant to geneticin were screened and confirmed by PCR assays.

### Site-directed mutagenesis and allelic exchanges

Allelic fragments with desired mutations, deletions, or insertions were generated using overlapping PCR, as previously described (*37*). For allelic exchanges at the native gene locus, the allelic fragments were connected to the N-terminal region of the hygromycin-resistance cassette through overlapping PCR. Fragments of approximately 1.0 kb downstream of the target regions were amplified and connected to the C-terminal region of the hygromycin-resistance cassette, also through overlapping PCR. Both fragments were co-transformed into PH-1 protoplasts. Transformants resistant to hygromycin were screened by PCR, and desired mutations were confirmed by DNA sequencing. Quantitative PCR was used to confirm that the transformants did not have any unintended integration of allelic fragments. For each mutation, at least two independent mutants with similar phenotypes were identified.

### Identification of repeat-induced point mutations in *FgTAD3*

To generate the tandemly repeated *FgTAD3 in situ*, the coding region of *FgTAD3* without the start codon was amplified and inserted in a reverse orientation before the promoter region (1832 bp upstream of the start codon) of *FgTAD3*. Two DNA fragments were generated using overlapping PCR. One fragment contained the upstream homologous fragment, the *FgTAD3* coding region, and the N-terminal region of the hygromycin-resistance cassette. The other fragment contained the C-terminal region of the hygromycin-resistance cassette and the downstream homologous fragment. These two DNA fragments were co-transformed into PH-1 protoplasts. Hygromycin-resistant transformants were screened and confirmed using PCR assays. Quantitative PCR was performed to ensure that unintended integration of the introduced fragments did not occur in the resulting TR-*FgTAD3* transformants.

For single ascospore isolation, ascospore cirrhi were collected from 10-dpf perithecia formed by the TR-*FgTAD3* transformants on carrot agar plates under a dissecting microscope and resuspended in 1 ml of sterile distilled water in a 1.5 ml centrifuge tube. The ascospore suspensions were then spread on 2% water agar, and ascospores with morphological defects were isolated using single spore isolation as described (*38*), and transferred onto PDA plates. The isolated single ascospores were then assayed for defects in growth and sexual reproduction. To identify mutations in the ascospore progeny, the endogenous *FgTAD3* gene was amplified from the DNA isolated from ascospore progeny and sequenced using Sanger sequencing.

### Rapid Amplification of 5’-Ends cDNA (5’RACE) and Reverse Transcription (RT)-PCR

Total RNA was extracted from hyphae or perithecia using the TRIzol reagent (Invitrogen, USA). The 5’RACE assays were performed using the FirstChoice^®^ RLM-RACE Kit (Invitrogen, USA) following the manufacturer’s instructions. In brief, mRNAs were converted into first-strand cDNA using reverse transcriptase Super Script^TM^ Ⅱ and a gene-specific primer 1 (GSP1). Following cDNA synthesis, the first-strand product was purified to remove unincorporated dNTPs and GSP1. Terminal deoxynucleotidyl transferase (TdT) was used to add homopolymeric tails to the 3’ ends of the cDNA. Tailed cDNA was then amplified by PCR using a gene-specific primer 2 (GSP2) downstream of GSP1 and an adapter primer that permitted amplification from the homopolymeric tail. The PCR products were detected by electrophoresis and connected to a T-vector for Sanger sequencing analysis.

For RT-PCR, cDNA synthesis was performed using RevertAid Master Mix (Thermo Scientific) following the manufacturer’s instructions. RT-PCR products were gel-purified and subjected to direct sequencing. Sanger sequencing traces were visualized using SnapGene Viewer 4.3 (https://www.snapgene.com/snapgene-viewer/) as described previously (*12*). Relative expression levels were assayed by quantitative real-time RT-PCR using the 2^−ΔΔCT^ method (*39*), with the actin gene serving as an internal control.

### Yeast two-hybrid (Y2H) assays

To detect protein interactions using yeast-two-hybrid (Y2H) assays, the open reading frames (ORFs) of genes and their mutant alleles were amplified and cloned into the pGADT7 prey or pGBKT7 bait vector (TaKaRa Bio, Tokyo, Japan). The resulting bait and prey constructs were transformed in pairs into yeast strain AH109 and assayed for growth on SD-His-Leu-Trp plates, as previously described (*40*). SD-Leu-Trp plates containing X-gal were used to assess the activity of LacZ β-galactosidase. SD-His-Leu-Trp or SD-His-Leu-Trp-Ade plates containing X-α-gal were used to evaluate the activity of Mel1 α-galactosidase.

### Western blotting

Total proteins were isolated and separated on 10% SDS-PAGE gels and transferred to nitrocellulose membranes, as described previously (*41*). The nitrocellulose membranes were incubated in 5% skim milk for 1 hour to avoid non-specific binding of antibodies. The proteins were then incubated with the anti-GFP (Abcam, ab290), anti-FLAG (Sigma-Aldrich, F31651), or anti-Tub2 (*42*) antibodies overnight at 4°C. After washing three times with 1×TBST (100 mM Tris-HCl, pH 7.5; 136 mM NaCl; 0.1% Tween-20), the proteins were incubated with corresponding secondary antibodies at room temperature for 1 hour. The non-specifically bound secondary antibodies were washed away with 1×TBST, and the proteins were incubated in a chemiluminescent substrate for 5-6 minutes before being detected by a chemiluminescence system.

### Co-immunoprecipitation (co-IP)

For co-IP to assay the interaction between FgTad2 and FgTad3, the P_RP27_-*FgTAD2*-3×FLAG and P_RP27_-*FgTAD3*-GFP fusion constructs were generated using the yeast gap-repair approach (*36*) and co-transformed into PH-1. Total proteins were isolated from 16 - h hyphae of the resulting transformants. The supernatant containing the target protein was incubated with anti-GFP affinity beads (SA070005, Smart-Lifesciences, China), and proteins were eluted using previously described methods (*41*). Western blots were performed on both total proteins and proteins eluted from the anti-GFP affinity beads using anti-FLAG (Sigma-Aldrich, F31651), anti-GFP (Abcam, ab290), or anti-Tub2 (*42*) antibodies for detection.

To investigate the interaction between Ame1 and FgTad3, fusion constructs of *AME1*-6×His and *FgTAD3*-Stag were created by inserting their ORFs into the MCS1 and MCS2 regions of the pRSFDuet-1 vector, respectively, which was then transformed into *E. coli* BL21. The resulting transformants were confirmed via PCR and western blot analysis. For co-IP assays, a 500 µL overnight bacterial culture was added to 50 mL of LB media and grown at 37°C until reaching an OD^600^ of 0.5-0.8. Recombinant protein expression was induced with isopropyl β-d-1-thiogalactopyranoside (IPTG) at a final concentration of 0.1 mM overnight at 20°C. Cells were lysed by sonication and the resulting clear lysate was harvested by centrifugation at 5000 rpm for 20 minutes, then transferred to a 4 mL tube. The supernatant containing the target protein was incubated with Ni-NTA agarose beads (Smart-Lifesciences, China) at 4°C overnight with gentle rocking for affinity purification. Western blots were performed on both total proteins and proteins eluted from the Ni-NTA beads using anti-His (CW0286M, CMBIO, China), anti-Stag (Cell Signaling Technology, 12774S), or anti-GAPDH (Sangon Biotech, D1100160200) antibodies.

### Assaying mRNA editing activities of the FgTad2-FgTad3-Ame1 complex in heterologous systems

To express *AME1*, M^4^-*FgTAD2*, and *FgTAD3* in yeast for assaying mRNA editing activities of the FgTad2-FgTad3-Ame1 complex, we amplified their ORFs and cloned them into pYES2, pYES-Leu, and pYES-Trp vectors, respectively, under the control of the *GAL1* promoter. The pYES-Leu and pYES-Trp vectors were derived from the pYES2 vector by digesting with *Nco*I and *Cla*I and replacing the Ura3 gene with Leu2 amplified from pGADT7 plasmid or Trp1 amplified from pGBKT7 plasmid. We co-transformed the resulting pYES2-*AME1*, pYES2-Leu-M^4^-*FgTAD2*, and pYES-Trp-*FgTAD3* vectors into yeast INVSc1 strain by heat shock. Transformants were confirmed by PCR and cultured in 10 mL 2% glucose SD-Ura-Leu-Trp liquid medium at 30°C for 24 hours. The cells were harvested by centrifugation and resuspended in 10 mL 2% galactose SD-Ura-Leu-Trp liquid medium at 30°C for 12 hours. We isolated total DNA and RNA from the cells and performed PCR and RT-PCR for *FgTAD3* and *AME1*. The purified PCR and RT-PCR products were subjected to direct Sanger sequencing for detecting editing events at the target A-to-I sites.

To assay mRNA editing activities of the FgTad2-FgTad3-Ame1 complex in bacteria, we cloned the codon-optimized ORFs of *AME1*, M^4^-*FgTAD2*, and *FgTAD3* into the pRSFDuet-1 plasmid under the control of both the T7 promoter and *lac* operon. We inserted *AME1* at the MCS1 region, while M^4^-*FgTAD2* and *FgTAD3* were sequentially inserted at the MCS2 region. We transformed the resulting vector into *E. coli* BL21 and confirmed the transformants by PCR. For control, we also transfected BL21 with the vector containing only M^4^-*FgTAD2* and *FgTAD3*. We cultured the transformants in LB media at 37°C and induced gene expression with isopropyl β-d-1-thiogalactopyranoside (IPTG) at a final concentration of 0.1 mM overnight at 20°C. We extracted total DNA and RNA and performed PCR and RT-PCR to assay the editing events in transcripts of *FgTAD3* by Sanger sequencing.

To assay mRNA editing activities of the FgTad2-FgTad3-Ame1 complex in human cell lines, we cloned the ORFs of *AME1*, M^4^-*FgTAD2*, and *FgTAD3* into the pCMV-Blank plasmid under the control of the CMV promoter. We used the generated vector to transfect the HEK 293T cell lines, respectively. For control, we also transfected the cell lines with the vectors containing M^4^-*FgTAD2* and *FgTAD3* or only *AME1*. After culturing at 37°C for 48 hours, we extracted total RNA and performed RT-PCR to assay the editing events in transcripts of *FgTAD3* and *AME1* by Sanger sequencing.

### Identification of post-translational modifications in FgTad3

We amplified the coding region of *FgTAD3* by PCR and cloned it into the vector pDL2 using the yeast gap-repair approach (*36*). The ORF of *FgTAD3* was fused with GFP and under the control of the RP27 promoter. The resulting *FgTAD3*-GFP fusion constructs were confirmed by Sanger sequencing analysis and transformed into PH-1. We confirmed hygromycin-resistant transformants expressing the fusion constructs by PCR and western blot analysis. Total proteins were isolated from hyphae harvested from 24-hour YEPD cultures and perithecia collected from mating plates at 7 dpf. We then incubated the proteins with anti-GFP affinity beads (SA070005, Smart-Lifesciences, China) at 4°C for 2.5 hours. The proteins were eluted from anti-GFP beads and detected by Coomassie blue staining. The FgTad3-GFP band cut out from the SDS-PAGE gel was digested with trypsin (Promega, USA) in gel at 37°C for 12 hours. Peptides were analyzed by the Thermo Scientific^™^ TSQ Quantum^™^ Access MAX triple quadrupole mass spectrometer, as in our previous reports (*37*).

### DNA-seq and strand-specific RNA-seq

DNA was extracted using the CTAB method. RNA was extracted and purified using the Eastep Super Total RNA Extraction Kit (Promega, USA). For eukaryotic RNA samples, poly (A)^+^ mRNA was enriched using magnetic beads with Oligo (dT), while for prokaryotic samples, rRNA was removed to enrich mRNA. DNA-seq and strand-specific RNA-seq libraries were constructed according to previous reports (*8, 11*) and sequenced on an Illumina NovaSeq 6000 system with 2×150-bp paired-end read mode at Novogene Bioinformatics Institute (Tianjin, China). Low-quality reads and reads containing adapters were removed by Trimmomatic (*43*) with default settings. The information regarding the DNA- and RNA-Seq data generated and utilized in this study has been listed in Table S6.

### RNA immunoprecipitation sequencing (RIP-seq)

We generated a strain expressing *FgTAD2*-3×FLAG at the native locus using the allelic exchange strategy described above. We collected 7-dpf perithecia produced by the *FgTAD2*-3×FLAG strain. RIP-seq assays were conducted with assistance from Wuhan IGENEBOOK Biotechnology (www.igenebook.com). In brief, the perithecia sample was cross-linked at 400 mj/cm2 at 4°C. Total proteins were then isolated, and 50 μL magnetic beads and 5 μg anti-Flag antibodies were mixed for 30 minutes at room temperature. The magnetic beads bound with antibodies were incubated with 900 μL total proteins overnight at 4°C for affinity purification. The FgTad2-3×FLAG proteins were eluted from magnetic beads for 30 minutes at 55°C. Input RNA and the RNA extracted from RIP eluate were used to construct sequencing libraries, respectively. The libraries were constructed with the NEBNext Ultra RNA Library Prep Kit following the manufacturer’s instruction and sequenced on an Illumina NovaSeq 6000 system with 2×150-bp paired-end read mode at Novogene Bioinformatics Institute (Tianjin, China).

### Analysis of RNA-seq and RIP-seq data

We obtained the reference genomes and annotation files of *F. graminearum* PH-1, *S. cerevisiae* R64-1-1, and *E. coli* BL21 from FgBase (http://fgbase.wheatscab.com/)(14), Ensembl Fungi, and NCBI genome database, respectively. After adapter trimming, the DNA-seq and RNA-seq reads were mapped to the reference genomes using Bowtie 2 v2.5.1 (*44*) and HISAT2 v2.2.1 (*45*) with the two-step model, respectively. Quality control of alignments was performed with Qualimap 2 (*46*). The number of reads aligned to each gene was calculated using featureCounts (*47*) and normalized by Reads Per Kilobase per Million mapped reads (RPKM) or TPM (Transcripts Per Million). For RIP-seq analysis, the normalized read density of each gene was calculated as Log_2_ (IP_RPKM/Input_RPKM+1). The genes expressed during sexual reproduction were divided into two groups: edited and unedited or high-edited and low-edited, using a dichotomous approach. The normalized read density was compared between groups. For A-to-I editing site identification, duplicated reads in the mapped RNA-seq BAM file were removed using the MarkDuplicates from Picard package v2.18.7 (http://broadinstitute.github.io/picard/). The resulting RNA-seq bam files were split into two bam files containing separated sense-strand and antisense-strand read alignments by BamTools v2.5.2 (*48*). A-to-I RNA editing sites were identified by REDItools v1.2 (*49*) using matched DNA-seq and RNA-seq data with an editing level cutoff value of 3%, as described previously (*50*). To facilitate comparison, an equivalent number of deduplicated mapped reads were randomly sampled. To exclude false positives, all A-to-G sites detected in samples with less than 100 sites were manually inspected using IGV (*51*).

The base preference surrounding the editing sites was visualized using WebLogo 3 (*52*). The secondary structures of 30-nt upstream and 30-nt downstream sequences of the editing or control sites were predicted using RNAFold in Viennarna v2.4.18 (*53*). We used forgi v2.0.3 (*54*) to perform statistics of secondary structure types. The random sampling of control sites with similar base preferences at −2 to +4 positions as edited sites was performed with the scripts developed in our previous study (https://github.com/wangqinhu/NC.edits) (*11*).

### Homology searches, phylogenetic analysis, and protein modeling

We identified gene homologs by performing a BLASTp search in the NCBI nr database and identified protein-conserved domains using NCBI CD-Search (https://www.ncbi.nlm.nih.gov/Structure/cdd/wrpsb.cgi). Gene orthologs were identified by orthAgogue (*55*) with default parameters. Multiple sequence alignments were performed with MUSCLE v5.1 (*56*) using default settings. The phylogenetic tree was constructed by IQ-TREE v2.2.2.7 (*57*) (-m MFP -T AUTO -B 1000) and visualized by iTOL (*58*). The protein complex structures of FgTad2-FgTad3-Ame1 and FgTad2-FgTad3 were predicted using AlphaFold-Multimer (*15*) on COSMIC2 (http://cosmic-cryoem.org/). The structures of FgTad2-FgTad3 and tRNA^Thr^_CGU_ were docked using PRIME v2.0.1 (*59*) with the model (PDB: 8AW3) of ADAT2/3 bound to tRNA from *Trypanosoma brucei* (*7*) as a template. All protein structures were visualized by ChimeraX v1.4 (*60*). We performed statistical significance tests with R (https://www.r-project.org/).

### Data availability

The data that support the findings of this study are available from the corresponding author upon request. Omics data generated in this study are accessible under the NCBI BioProject accession: PRJNA1020378.

