## Supplemental Figure 1 to 6 for "Unveiling the A-to-I mRNA editing machinery and its regulation and evolution in fungi"

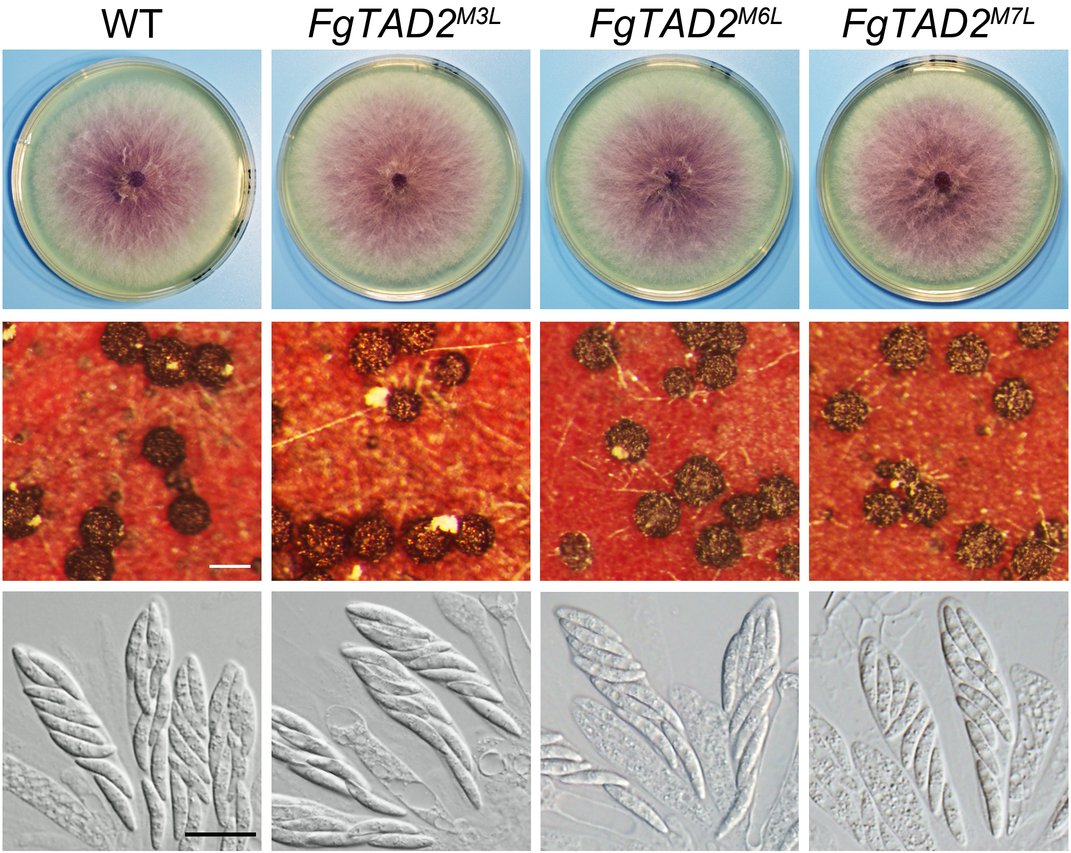


**Fig. S1. Normal vegetative growth and sexual development in *FgTAD2*$^{M3L}$, *Fg-TAD2*$^{M6L}$, and *FgTAD2*$^{M7L}$ mutants.** White bar = 0.2 mm; black bar = 20 μm.

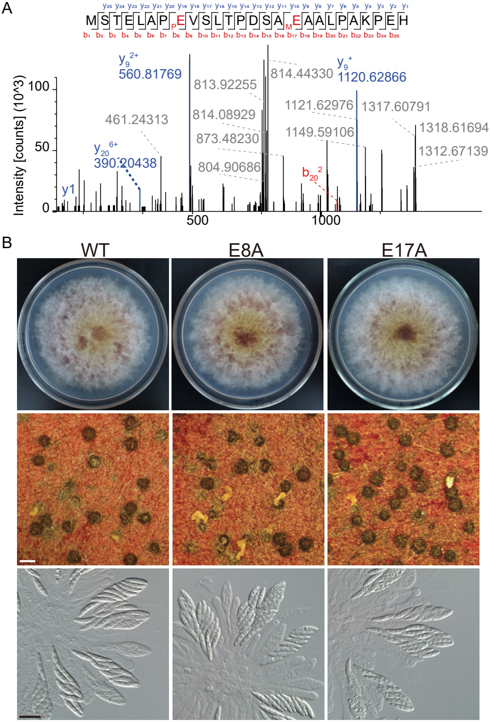


**Fig. S2. Normal vegetative growth and sexual development in the post-translational modification-deficient mutants $FgTAD3^{E8A}$ and $FgTAD3^{E17A}$.** (A) MS/MS spectrum of the peptide containing the phosphorylated ($pE^{8}$) and methylated ($mE^{17}$) residues. (B) Colony morphology, perithecium formation, and asci/ascospores morphology of the modification-deficient mutants $FgTAD3^{E8A}$ and $FgTAD3^{E17A}$. White bar = 0.2 mm; black bar = 20 μm.

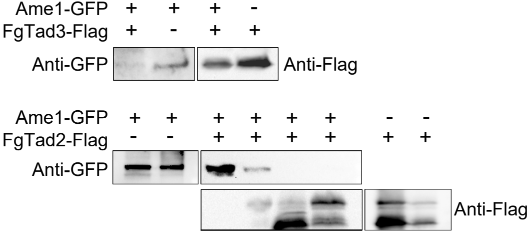


**Fig. S3. Western blots showing that the expression of Ame1-GFP was suppressed when coexpressed with FgTad2-FLAG or FgTad3-FLAG.**
